## Supplemental Files 1-7 for "Is male dimorphism under sexual selection in humans? A meta-analysis"

Supplementary File 1

*Effect size conversion formulas*

| Kendall’s tau | r = sin (.5 πτ) (Kendall, 1970) |
| --- | --- |
| Spearman's rho | not converted |
| t | rYλ = √(t^2^ / (t^2^ + df)) Online converter: <https://www.uccs.edu/lbecker/> |
| Odds ratio | Online converter: http://escal.site/ |
| Unstandardized regression coefficient (B) | β = (S.D. of predictor/S.D. of outcome) × B; r = .98β + .05λ (λ = 1 when β is nonnegative and 0 when β is negative) |
| Standardized regression coefficient (β) | r = .98β + .05λ (λ = 1 when β is nonnegative and 0 when β is negative; Peterson & Brown, 2005) |

Supplementary File 2A

*General coding decisions*

| Sexual orientation | Coded as non-heterosexual sample if the sample was mixed but predominantly heterosexual. |
| --- | --- |
| Samples or subsamples comprising only fathers and/or married individuals | Coded as heterosexual unless otherwise specified, since they had reproduced/married heterosexually. |
| Student samples with a mean age ≤ 20 | Coded as non-fathers. |
| Sample contained ≥ 50% students | Coded as a student sample. |
| Sample contained both students and non-students but the proportion of students/non-students was not mentioned | Coded as a non-student sample. |
| Age | Considered an essential control for all outcome variables except age at first sexual intercourse/encounter and age at the birth of the first child, unless all participants were the same age, and for mating attitude measures. |
| Ethnicity | Coded as ‘white’ if ≥ 75% of sample was white. |
| Marriage system | Coded as polygynous if polygyny was permitted in population, even if rare. |
| Online samples | Coded as low fertility and monogamous. |
| Cut off point for high versus low fertility | 3.0 children/woman (in sample or population at the time of sampling). |
| Extreme outliers | Were included when possible, as outliers were expected. |
| Analyses of relevant relationships were included in paper, but authors had submitted results/raw data to us (e.g. results for men only, controlling for age etc.). | Coded as published results. |
| Preprints | Coded as non-published and non-peer-reviewed, unless the paper was later accepted for publication in which case it was updated as published and peer-reviewed. |
| Paper contained both zero-order correlations and multiple regression coefficients | We chose the regression coefficient if the multiple regression included relevant control variables (such as age), and the correlation coefficient if the multiple regression included irrelevant control variables. |
| Effect sizes given as Spearman’s rho | Were not converted; however, were coded as converted for moderation analyses, because it was not given as *r* and therefore considered to be an estimate. |
| Number of children in industrialized populations | Coded as children born (rather than surviving children) unless otherwise specified. (In naturally fertile populations, it is typically spelled out whether measures refer to number of children born vs number of surviving children.) |
| Dataset on age at first sexual intercourse contained virgins | Current age was used. |
| Testosterone studies where the authors only included samples that were clear | Coded as having controlled for blood contamination. |
| Muscularity measures | When other-rated, adiposity should be controlled for and was thus considered a necessary control; when own-rated, adiposity was not considered a necessary control, since people should be able to assess their own amount of muscle/adiposity. |
| Handgrip strength | Moderator ‘number of measurements’ was coded as number of measurements per hand, not in total. |

| Supplementary File 2B  *Study-specific coding decisions* | |
| --- | --- |
| Authors | Decisions |
| Alvergne et al., 2009 | We assumed N=53 (married fathers only) as *p*-value does not add up if whole sample of married and non-married was analyzed together. It also makes sense to only analyze married men as they were the only ones who were able to reproduce. Not explicit in papers which variables were transformed to normality. |
| Apicella, 2014 | We excluded DVs >1 spouse in lifetime (considered redundant) and number of offspring born (effect size is the same for another variable but *p*-value differs - N is not stated, suggesting that either the effect size is not correct or N is considerably smaller). N for some analyses is not given, we assumed it was 51 as given for one of the analyses. They classified predictor as strength, but we re-coded it as a composite measure of muscle mass and handgrip strength as that is what it was (predictor was therefore not classified as either muscle mass or strength for moderation analyses). Some relationships reported in other papers as well: Smith, Olkhov, Puts & Apicella (2017) reported muscle mass/strength – reproductive success and offspring number; we kept results from this paper as it controlled for age, with the exclusion of offspring number for reason given above. |
| Apicella et al., 2007 | Some relationships reported in other papers as well: Smith, Olkhov, Puts & Apicella (2017) reported f0 - reproductive success and offspring number; we kept results from this paper as it controlled for age. |
| Arnocky et al., 2018 | Paper also included fWHR-lower - SOI-R and lower face/face height - SOI-R but effect sizes not reported separately for men and women so not included. |
| Atkinson et al., 2012 | Paper included both DVs number of living children and genetic vector; the latter calculated as 1*(number of living children) + ½*(number of living grandchildren). Considered redundant to include both, and to be consistent with other measures, we included number of living children. |
| Boothroyd et al., 2017 | Agta sample: photographs taken from front or ¾ degree angle were coded as not frontal photographs. |
| Charles & Alexander, 2011 | We excluded SOI (Clark, 2004) as it is redundant to SOI and SOI is the commonly used measure. Sample assumed to be non-fathers. |
| Falcon, 2016 | We used average 2D:4D rather than R2D:4D/L2D:4D due to bigger sample size and we could not rule out the possibility of overlapping samples. |
| Farrelly et al., 2015 | All participants were heterosexual (information provided by author). Author re-ran analyses based on whole sample and provided results which we used, so the results do not exactly match results reported in paper (in the paper the authors had omitted a few participants due to incomplete relationship information). |
| Frederick & Jenkins, 2015 | The paper also included dichotomous variables: more than 5 sex partners and more than 14 sex partners. We did not include those (considered redundant) and instead only used the continuous variable number of sex partners. |
| Gallup et al., 2007 | The paper included both SHR circumference and breadth; we only included circumference to keep it consistent with other results. |
| Genovese, 2008 | In the paper, the relationship between HGS and height was reported, but the paper did not include N or information about whether age was controlled for, so first author re-ran analyses. For mesomorphy - offspring number, we assumed N=181 as first author could only find (reliable) data on offspring number for 181 participants in the primary data source. |
| Gettler et al., 2019 | Fertility in The Philippines has now dropped below 3.0 children/woman but was above in 2009 when data was collected (according to https://data.worldbank.org/indicator/SP.DYN.TFRT.IN?locations=PH), therefore coded as a high fertility sample. |
| Hartl et al., 1982 | HGS - offspring number and height - offspring number were also analyzed and reported by Genovese (2008), but that paper did not include N and it was not clear whether age had been controlled for, so first author re-ran analyses. Participants with clearly incomplete or inaccurate family histories were excluded. In cases where family history was clear up to a certain point, or the participant had died, their age at that point was used. Thus, some of these relationships are reported as non-peer reviewed and some as peer-reviewed (the latter in the case where the relationship was reported by Genovese). We did not include general strength, as it was assessed subjectively. |
| Hoppler, Walther et al., 2018 | Ninety-seven percent of sample was white (mentioned in other paper on the same sample). |
| Hönekopp et al., 2007 | Same sample as in Hönekopp et al., 2006, who reported that 80% of the sample were students. |
| Kirchengast, 2000 | Judged to be the same sample as Winkler and Kirchengast, 1994. |
| Kirchengast & Winkler, 1995 | Mean number of children in sample: 1.1 in Rundu and 1.8 in rural areas; however, age range of sample was 18-39 and mean age = 26 so it is young sample with non-completed reproductive histories, therefore coded as a high fertility sample. |
| Klimas et al., 2019 | Ninety-seven percent of sample was white (mentioned in other paper on same sample). Only included men without sexual dysfunction. Excluded participants who had had bleeding or injuries in the mouth in the last few days before testing (clear from other paper on same sample), so we considered blood contamination of saliva sample controlled for. |
| Little et al., 1989 | Population mean and S.D. for height stated in another paper (by the same author); we used that to convert effect sizes. |
| Loehr & O'Hara, 2013 | We assumed it was primarily a white sample. |
| Longman et al., 2018 | Given that this was a young British student sample, we assumed that they were all non-fathers. Baseline testosterone was, in a sense, anticipatory, but we included this paper since effect sizes in this study did not differ substantially from effect sizes in other studies. |
| Lukaszewski et al., 2014 | The first author ran analyses on openly available data. Some extreme outliers in one of the samples (chest strength around ~2, which seems incorrect). However, it made no difference to the results whether they were removed or kept, so we kept them in. |
| Marczak et al., 2018 | 2D:4D was measured directly as well as from digital photos. Not explicit whether that meant that measurement method varied between participants. As we could not be sure that all participants had been measured directly, we coded this as hand scans. |
| Mosing et al., 2015 | Published paper but author submitted results to us. Twin sample; only unrelated individuals included in this sample. We coded it as a heterosexual sample (in the paper, gay participants were excluded), and as a predominantly white sample. |
| Mueller & Mazur, 1997 | N was not explicit in the results, but they stated that 337 participants replied so we assumed N=337. Sample was born 1923-1929 so should have completed most of their reproduction by 1965 when the fertility rate dropped below 3 in the U.S.; therefore coded as a high fertility, industrialized sample. |
| Nagelkerke et al., 2006 | Author sent us raw data. For age at first sexual intercourse, we set cut off at 12 (there were data points <12) as we deemed it unlikely that participants had had sexual onset prior to puberty. |
| Nettle, 2002 | This sample's parents were analyzed in Krzyzanowska et al. (2015) but as the parents reproduced separately and at a different time point compared to this sample, we considered them to be separate samples. |
| Pawlowski et al., 2008 | We assumed this sample was heterosexual. |
| Pawlowski et al., 2000 | Author sent us the results. Coded as a high fertility, industrialized sample. |
| Polo et al., 2019 | The paper also included skeletal muscle mass but this measure was extremely highly correlated with upper-body fat free mass (FFM: *r* = .96, n = 206, *p* < .001) so it was considered redundant to include both predictors, and to be consistent with other studies we kept upper-body FFM. Sample consisted of heterosexual students and non-students: we did not know the proportion of students versus non-students, so we coded it as a non-student sample. Results were considered published as these relationships were reported in paper, although the paper included a type of analyses that we could not use and the authors therefore submitted *r*. |
| Puts et al., 2015 | We assumed the sample were non-fathers. For one sample, testosterone was sampled by saliva between 9AM-1.30PM; we coded that as AM testosterone. |
| Puts et al., 2006 | We judged the sample to be the same one as Putz et al. (2004) and Hodges-Simeon et al. (2011). |
| Putz et al., 2004 | We judged the sample in study 1 to be the same one as Puts et al. (2006) and Hodges-Simeon et al. (2011). |
| Rahman et al., 2005 | Proportion of students vs non-students not clear, so coded as unknown low fertility sample. |
| Rosenfield et al., 2020 | Coded as heterosexual sample as all participants had been married (heterosexually) at some point. |
| Scott & Bajema, 1982 | The paper reported both zero-order correlations and partial correlations controlling for ethnicity. We used zero-order correlations because the sample was from the same group, even if their ethnicities differed. Sample was born 1912-1918 and should therefore have largely completed reproduction by 1965 when the fertility rate dropped below 3 in the U.S.; therefore coded as a high fertility, industrialized sample. |
| Sim & Chun, 2016 | We judged the sample to be the same one as Sim (2013). SHR also reported in Sim (2013); we therefore excluded that paper. |
| Smith et al., 2017 | Same sample/analyses as reported in Apicella et al. (2007) and Apicella (2014); kept those papers as those analyses controlled for age, with the exception of muscle mass/strength - offspring number (also given in Apicella, 2014), as in the latter, N was not explicit. |
| Steiner et al., 2011 | Paper also included SOI and extrapair sexual interest (EPSI), but SOI and EPSI were measured after viewing a video which, for some participants, had sexual content, and those variables were therefore measured after manipulation so we did not include them. Sample consisted of 90% exclusively heterosexual participants, and 10% heterosexual but incidentally gay participants; coded as a heterosexual sample. |
| Stern et al., 2020 | Information about ethnicity not available in paper; however, it was given in other paper on same sample (Kandrik et al., 2016) in which it was reported that 91% of a subsample was white. This sample was therefore coded as predominantly white. |
| Strong et al., 2014 | We excluded 2D:4D following recommendation from author, due to potential measurement issues. |
| Suire et al., 2018 | Baseline recording consisted of just a short utterance repeated after the experimenter, and author therefore suggested to use one of the other recordings (courtship and competition) instead. Results did not show any substantial difference between the recordings, however, so we used baseline to be consistent with other studies. Data set contained virgins: current age set as age at first sexual intercourse, as is commonly done. |
| Tao & Yin, 2016 | Offspring number: S.D. of the mean for the whole sample not given but varied between 1.1-1.3 for each of the three samples so we used that to convert effect sizes (which value was used did not affect the results in any case). We used whole sample rather than the three sub-samples, coded as low fertility. |
| van Anders et al., 2007 | Sampled from a monogamous society, but some of the samples were polyamorous, therefore coded as non-monogamous. |
| van Dongen & Sprengers, 2012 | Sample not specified but we assumed it was from a low fertility, monogamous population. |
| Varella et al., 2014 | Assumed N=80 for all non-significant relationships where N was not specified, as stated elsewhere in the paper. We used results for the two samples combined, not the sub-samples. |
| von Rueden et al., 2010 | DVs not normally distributed but could be transformed to near-normality. |
| Walther et al., 2016 | Ninety-seven percent of sample was white (mentioned in other paper on same sample). |
| Walther et al., 2017b | Ninety-seven percent of sample was white (mentioned in other paper on same sample). |
| Walther et al., 2017c | Ninety-seven percent of sample was white (mentioned in other paper on same sample). |
| Weeden & Sabini, 2007 | Paper also included Sociosexuality measure; however, as we did not know the response scale or direction of responses, we excluded it. |
| Winkler & Kirchengast, 1994 | Judged to be the same sample as Kirchengast, 2000. |

*Note*. 2D:4D = 2^nd^ to 4^th^ finger (digit) ratio; f0 = fundamental frequency, i.e. voice pitch; fWHR = facial width-to-height ratio; HGS = handgrip strength; SHR = shoulder-to-hip ratio; SOI = Sociosexual Orientation Inventory; SOI-R = the Revised Sociosexual Orientation Inventory.

| Supplementary File 3A  *General moderators: for all predictors* | |
| --- | --- |
| Moderator | Description |
| Domain type  Mating measure type | Mating vs reproductive domain.  Mating attitudes (e.g. preferences for short-term relationships/casual sex) vs mating behaviors (e.g. number of sexual partners, age at first sexual intercourse) |
| Reproductive measure type | Fertility (i.e. number of children/grandchildren, age at the birth of the first child) vs reproductive success (i.e. number of surviving children/grandchildren). |
| Sample type | Low fertility samples (i.e. <3.0 children/woman within sample/population at the time of sampling) vs high fertility samples. The latter is considered to correspond to naturally fertile populations. |
| Low fertility samples | Predominantly student samples (i.e. ≥ 50% students) vs mixed/non-student/unknown samples. |
| High fertility samples | Traditional vs industrialized samples. |
| Ethnicity | Predominantly white (i.e. ≥ 75% of sample) vs mixed/non-white /unknown. |
| Marriage system | Monogamy vs non-monogamy/unknown. |
| Publication status | Published vs non-published results. We favored publication status of the relevant *results* rather than of the *paper*, since we retrieved many of our effects from published studies where the key relationship had not been analyzed/was not the focus of the paper. |
| Peer-review | Peer-reviewed vs not peer-reviewed study. |
| Sexual orientation | Heterosexual sample vs gay/mixed/unknown sexual orientation. |
| Normality-transformed variables | Non-transformed vs transformed variables, i.e. whether skewed variables (skew is very common for some of the variables, such as number of sexual partners) had been e.g. log-transformed to normality. |
| Converted effect size | Non-converted vs converted effect sizes, i.e. whether effect size was given as Pearson’s *r* or whether we had used a formula to convert it. The latter results in an estimate of *r*. |
| Age control | Age controlled for in analyses vs not controlled for. We considered age an essential control for all analyses except *i*. where all participants belonged to the same age group, *ii.* for the variables sexual onset/reproductive onset, and *iii.* mating attitudes. |
| Non-relevant controls | No non-relevant vs non-relevant controls included in the analyses. For example, analyses with several non-relevant predictors may produce weaker associations compared to e.g. bivariate correlations with just the relevant predictor and outcome variables. |
| *Note.* We were constrained by information made available in papers. The levels of moderators should therefore be considered to reflect where we knew for certain that a moderator e.g. had been controlled for vs where we could not be certain. For several of our potential moderators, such as the moderators we had selected for voice pitch, not enough papers mentioned having controlled for them and we were therefore unable to analyze those moderators. Additionally, we often did not have enough observations on each level of the moderator variable to be able to run those analyses. This lack of power also prevented us from analyzing combined effects of several moderators; we therefore analyzed moderators one by one. Moderators were coded into two levels wherever possible, as otherwise we would often have had too few observations/level to be able to run the analysis. It should also be noted that some moderators are likely confounded; for example, non-monogamous populations are almost always high fertility, traditional populations. | |

Supplementary File 3B

| *Facial masculinity moderators* | |
| --- | --- |
| Moderator | Description |
| Measurement type | Objectively measured masculinity (using geometric morphometric analyses) vs observer-rated masculinity vs fWHR (i.e. facial width-to-height ratio). |
| Standardization of photographs | Photographs taken under standardized vs not standardized/semi-standardized/unknown conditions. |
| Angle of photographs | Front-facing vs not front-facing/unknown angle of photographs. |
| Masked photographs | Masked vs not masked photographs/unknown. Only coded for rated facial masculinity. |
| Adiposity | Adiposity/body mass index (BMI) controlled for vs not controlled for/unknown. Only coded for rated facial masculinity. |
| Color vs black & white photographs | Color vs black & white photographs/unknown. Only coded for rated facial masculinity. |
| Facial expression | Neutral vs smiling/mixed/unknown facial expressions. Only coded for rated facial masculinity. |
| Facial hair | Clean-shaven vs not clean-shaven/mixed/unknown. Only coded for rated facial masculinity. |
| *Note.* Moderators were coded into two levels wherever possible, as otherwise we would often have had too few observations/level to be able to run the analysis. | |

Supplementary File 3C

| *Body masculinity moderators* | |
| --- | --- |
| Moderator | Description |
| Number of measurements | Only coded for measured body masculinity. Typically referred to repeat measurements but in some cases, different measurements were used. |
| Adiposity | Adiposity/BMI controlled for vs not controlled for/unknown. Only coded for rated body masculinity. |
| Measurement type | Measured vs observer- or own-rated body masculinity. Measured body masculinity included e.g. strength, circumference of shoulder-to-hip ratio, and bioelectrical measurement of fat-free mass. |
| Body masculinity type | Strength vs body shape vs muscle mass. Strength was typically assessed through measured handgrip strength. Body shape included body measurements (see measurement type above) and rated body masculinity. Muscle mass was measured (see above) or rated. |
| *Note.* Moderators were coded into two levels wherever possible, as otherwise we would often have had too few observations/level to be able to run the analysis. | |

Supplementary File 3D

| *2D:4D moderators* | |
| --- | --- |
| Moderator | Description |
| Measurement type | Measured directly vs measured from hand scans/photographs vs self-reported vs unknown. |
| Number of measurements | Only coded for experimenter-measured (directly or from hand scans). |
| Finger injuries | Controlled for vs not controlled for. |
| Left vs right | Left vs right hand 2D:4D. 2D:4D dimorphism is typically claimed to be more pronounced in the right hand (Hönekopp et al., 2006). |
| *Note.* Moderators were coded into two levels wherever possible, as otherwise we would often have had too few observations/level to be able to run the analysis. | |

Supplementary File 3E

| *Voice pitch moderators* | |
| --- | --- |
| Moderator | Description |
| Sex of experimenter | Female vs male vs unknown. |
| Illness | Illnesses (colds etc. that could influence voice pitch) controlled for/excluded vs not. |
| Smoker | Smoking participants controlled for/excluded vs not. |
| Condition | Baseline vs courtship cs competitive type of recording. |
| *Note.* Based on information available in the papers, none of these potential moderators had been controlled for in any of the studies. | |

Supplementary File 3F

| *Height moderators* | |
| --- | --- |
| Moderator | Description |
| Measurement type | Experimenter-measured vs self-reported vs unknown. |
| Number of measurements | Only coded for experimenter-measured height. |
| *Note.* Moderators were coded into two levels wherever possible, as otherwise we would often have had too few observations/level to be able to run the analysis. | |

Supplementary File 3G

| *Testosterone levels moderators* | |
| --- | --- |
| Moderator | Description |
| How assayed | Assayed from blood vs saliva vs unknown. |
| Time of day | Assayed in the AM vs PM vs unknown. |
| Blood contamination | Checked for vs not checked for/unknown. Only coded for saliva assayed T levels. |
| Fatherhood | Non-fathers vs fathers vs mixed/unknown |
| Relationship status | Married/in committed relationship vs single vs mixed/unknown. |
| *Note.* Moderators were coded into two levels wherever possible, as otherwise we would often have had too few observations/level to be able to run the analysis. | |

Supplementary File 4A

| *Facial masculinity: moderation analyses. The intercept shows the 'simple effect' for the reference category (specified) and the moderator effect shows the change in effect size for that category relative to the reference category. Moderators are bolded if significant after controlling for multiple comparisons, as indicated by computation of q-values. The full list of q-values can be found in Supplementary File 7.* | | | | | | |
| --- | --- | --- | --- | --- | --- | --- |
| Mating vs reproductive domain \| \| \| \| \| \|; \| \| B \| SE \| [95% CI] \| *z* \| *p*; Domain type \| Intercept (mating domain) \| 0.088 \| 0.042 \| 0.007, 0.169 \| 2.120 \| .034; \| Reproductive domain \| 0.006 \| 0.072 \| -0.135, 0.148 \| 0.089 \| .929 | | | | | | |
| Mating domain (MAT), mating behaviors & mating attitudes | | | | | | |
|  |  | B | SE | [95% CI] | *z* | *p* |
| MAT measure type | Intercept (MAT behaviors) | 0.047 | 0.041 | -0.033, 0.128 | 1.150 | .250 |
|  | MAT attitudes | 0.038 | 0.061 | -0.082, 0.159 | 0.622 | .534 |
| Ethnicity | Intercept (predominantly white) | 0.114 | 0.057 | 0.003, 0.225 | 2.007 | .045 |
|  | Mixed/other/unknown | -0.075 | 0.084 | -0.241, 0.090 | -0.892 | .373 |
| Publication status | Intercept (published results) | 0.046 | 0.058 | -0.067, 0.159 | 0.801 | .423 |
|  | Non-published results | 0.092 | 0.093 | -0.090, 0.273 | 0.990 | .322 |
| Publication status: MAT behaviors | Intercept (published results) | 0.016 | 0.056 | -0.094, 0.126 | 0.281 | .778 |
|  | Non-published results | 0.023 | 0.098 | -0.169, 0.215 | 0.233 | .816 |
| Sexual orientation | Intercept (heterosexual sample) | 0.044 | 0.055 | -0.063, 0.151 | 0.806 | .420 |
|  | Gay/mixed/unknown | 0.105 | 0.092 | -0.075, 0.285 | 1.145 | .252 |
| Age control | Intercept (age controlled for) | 0.065 | 0.060 | -0.052, 0.181 | 1.086 | .277 |
|  | Age not controlled for | 0.038 | 0.098 | -0.154, 0.230 | 0.387 | .699 |
| Age control: MAT behaviors | Intercept (age controlled for) | 0.017 | 0.052 | -0.086, 0.119 | 0.318 | .750 |
|  | Age not controlled for | 0.024 | 0.100 | -0.172, 0.220 | 0.241 | .810 |
| Measurement type | Intercept (measured) | 0.105 | 0.058 | -0.008, 0.219 | 1.825 | .068 |
|  | Rated | -0.009 | 0.079 | -0.162, 0.146 | -0.108 | .914 |
|  | fWHR: *s* =2 |  |  |  |  |  |
| Other moderators with too few *k*/*s*: | | | | | | |
| Sample type: low vs high fertility; Low fertility samples: predominantly students vs non-students; High fertility samples: traditional vs industrialized; Marriage system: monogamy vs non-monogamy; Peer-reviewed vs not peer-reviewed; Normality-transformed variables; Converted effect size; Non-relevant controls, Standardization of photographs; Angle of photographs; Masked photographs; Adiposity; Color vs black & white photographs; Facial expression; Facial hair. | | | | | | |
| Reproductive domain (REP), fertility & reproductive success | | | | | | |
| Moderators with too few *k*/*s*: | | | | | | |
| REP measure type: reproductive success vs fertility; Sample type: low vs high fertility; Low fertility samples: predominantly students vs non-students; High fertility samples: traditional vs industrialized; Ethnicity: predominantly white vs not; Marriage system: monogamy vs non-monogamy; Publication status: published results; Peer-reviewed; Sexual orientation: heterosexual sample vs gay/mixed/unknown; Normality-transformed variables; Converted effect size; Age controlled for; Non-relevant controls, Measurement type: measured vs rated vs fWHR; Standardization of photographs; Angle of photographs; Masked photographs; Adiposity; Color vs black & white photographs; Facial expression; Facial hair. | | | | | | |
| *Note.* *k* = number of observations, MAT = mating, REP = reproductive, *s* = number of samples. Moderation analyses were only run where each level of the moderator included observations from at least two studies and three independent samples. Analyses were run on the mating measures mating behaviors and mating attitudes, and the reproductive measures fertility and reproductive success when there were enough observations to do so. | | | | | | |

Supplementary File 4B

| *Body* *masculinity: moderation analyses. The intercept shows the 'simple effect' for the reference category (specified) and the moderator effect shows the change in effect size for that category relative to the reference category. Moderators are bolded if significant after controlling for multiple comparisons, as indicated by computation of q-values. The full list of q-values can be found in Supplementary File 7.* | | | | | | |
| --- | --- | --- | --- | --- | --- | --- |
| Mating vs reproductive domain \| \| \| \| \| \|; \| \| B \| SE \| [95% CI] \| *z* \| *p*; Domain type \| Intercept (mating domain) \| 0.132 \| 0.021 \| 0.092, 0.173 \| 6.414 \| <.001; \| Reproductive domain \| 0.019 \| 0.042 \| -0.064, 0.102 \| 0.439 \| .661 | | | | | | |
| Mating domain (MAT), mating behaviors & mating attitudes | | | | | | |
|  |  | B | SE | [95% CI] | *z* | *p* |
| MAT measure type | Intercept (MAT behaviors) | 0.139 | 0.022 | 0.096, 0.181 | 6.382 | <.001 |
|  | MAT attitudes | -0.024 | 0.031 | -0.085, 0.037 | -0.773 | .440 |
| Sample type | Intercept (low fertility) | 0.136 | 0.023 | 0.091, 0.181 | 5.878 | <.001 |
|  | High fertility | -0.024 | 0.078 | -0.177, 0.129 | -0.307 | .759 |
| Sample type: MAT behaviors | Intercept (low fertility) | 0.147 | 0.024 | 0.099, 0.194 | 6.019 | <.001 |
|  | High fertility | -0.035 | 0.078 | -0.188, 0.119 | -0.443 | .658 |
| Low fertility sample | Intercept (predominantly students) | 0.157 | 0.023 | 0.111, 0.203 | 6.699 | <.001 |
|  | Non-students/ mixed/unknown | -0.118 | 0.051 | -0.218, -0.019 | -2.336 | .020 |
| Low fertility sample: MAT behaviors | Intercept (predominantly students) | 0.172 | 0.024 | 0.125, 0.218 | 7.285 | <.001 |
|  | **Non-students/ mixed/unknown** | **-0.128** | **0.049** | **-0.224, -0.033** | **-2.632** | **.009** |
| Ethnicity | Intercept (predominantly white) | 0.116 | 0.044 | 0.030, 0.203 | 2.643 | .008 |
|  | Mixed/other/unknown | 0.024 | 0.051 | -0.076, 0.124 | 0.464 | .643 |
| Ethnicity: MAT behaviors | Intercept (predominantly white) | 0.116 | 0.044 | 0.031, 0.202 | 2.661 | .008 |
|  | Mixed/other/unknown | 0.038 | 0.052 | -0.064, 0.139 | 0.728 | .467 |
| Marriage system | Intercept (monogamy) | 0.139 | 0.022 | 0.095, 0.182 | 6.230 | <.001 |
|  | Non-monogamy | -0.095 | 0.096 | -0.283, 0.093 | -0.991 | .322 |
| Marriage system: MAT behaviors | Intercept (monogamy) | 0.149 | 0.023 | 0.104, 0.195 | 6.422 | <.001 |
|  | Non-monogamy | -0.106 | 0.096 | -0.294, 0.082 | -1.104 | .270 |
| Publication status | Intercept (published results) | 0.167 | 0.026 | 0.117, 0.218 | 6.470 | <.001 |
|  | Non-published results | -0.086 | 0.039 | -0.163, -0.009 | -2.181 | .029 |
| Publication status: MAT attitudes | Intercept (published results) | 0.099 | 0.044 | 0.013, 0.184 | 2.251 | .024 |
|  | Non-published results | -0.079 | 0.081 | -0.237, 0.079 | -0.975 | .330 |
| Publication status: MAT behaviors | Intercept (published results) | 0.177 | 0.028 | 0.123, 0.231 | 6.402 | <.001 |
|  | Non-published results | -0.087 | 0.044 | -0.172, -0.001 | -1.978 | .048 |
| Peer-review | Intercept (peer-reviewed) | 0.136 | 0.024 | 0.088, 0.184 | 5.576 | <.001 |
|  | Not peer-reviewed | -0.012 | 0.058 | -0.126, 0.102 | -0.204 | .838 |
| Peer-review: MAT behaviors | Intercept (peer-reviewed) | 0.142 | 0.025 | 0.093, 0.192 | 5.636 | <.001 |
|  | Not peer-reviewed | 0.005 | 0.063 | -0.118, 0.128 | 0.085 | .933 |
| Sexual orientation | Intercept (heterosexual sample) | 0.177 | 0.030 | 0.118, 0.235 | 5.948 | <.001 |
|  | Gay/mixed/unknown | -0.085 | 0.041 | -0.165, -0.006 | -2.098 | .036 |
| Sexual orientation: MAT attitudes | Intercept (heterosexual sample) | 0.045 | 0.057 | -0.067, 0.157 | 0.781 | .435 |
|  | Gay/mixed/unknown | 0.062 | 0.077 | -0.089, 0.212 | 0.804 | .421 |
| Sexual orientation: MAT behaviors | Intercept (heterosexual sample) | 0.188 | 0.031 | 0.127, 0.249 | 6.069 | <.001 |
|  | Gay/mixed/unknown | -0.088 | 0.042 | -0.171, -0.006 | -2.091 | .037 |
| Normality-transformed variables | Intercept (non-transformed variables) | 0.137 | 0.025 | 0.088, 0.185 | 5.523 | <.001 |
|  | Transformed variables | 0.038 | 0.047 | -0.054, 0.129 | 0.810 | .418 |
| Normality-transformed variables: MAT behaviors | Intercept (non-transformed variables) | 0.049 | 0.037 | -0.024, 0.122 | 1.321 | .186 |
|  | **Transformed variables** | **0.165** | **0.054** | **0.060, 0.270** | **3.091** | **.002** |
| Converted effect size | Intercept (not converted) | 0.144 | 0.026 | 0.093, 0.194 | 5.587 | <.001 |
|  | Converted | 0.003 | 0.063 | -0.121, 0.128 | 0.053 | .958 |
| Converted effect size: MAT behaviors | Intercept (not converted) | 0.152 | 0.028 | 0.098, 0.206 | 5.508 | <.001 |
|  | Converted | -0.005 | 0.065 | -0.133, 0.123 | -0.076 | .940 |
| Age control | Intercept (age controlled for) | 0.098 | 0.031 | 0.037, 0.158 | 3.147 | .002 |
|  | **Age not controlled for** | **0.103** | **0.042** | **0.020, 0.186** | **2.441** | **.015** |
| Age control: MAT behaviors | Intercept (age controlled for) | 0.107 | 0.033 | 0.043, 0.171 | 3.277 | .001 |
|  | Age not controlled for | 0.096 | 0.045 | 0.009, 0.183 | 2.153 | .031 |
| Number of measurements | Intercept (unknown number of measurements) | 0.076 | 0.041 | -0.005, 0.156 | 1.851 | .064 |
|  | 2 measurements | 0.046 | 0.047 | -0.046, 0.137 | 0.974 | .330 |
|  | 3 measurements | 0.126 | 0.062 | 0.004, 0.247 | 2.025 | .043 |
|  | 1 measurement: *s* = 1 |  |  |  |  |  |
| Number of measurements: MAT attitudes | Intercept (unknown number of measurements) | 0.117 | 0.054 | 0.010, 0.223 | 2.144 | .032 |
|  | 2 measurements | -0.070 | 0.071 | -0.210, 0.069 | -0.987 | .324 |
|  | 1 measurement: *s* = 1 |  |  |  |  |  |
|  | 3 measurements: *s* = 0 |  |  |  |  |  |
| Number of measurements: MAT behaviors | Intercept (unknown number of measurements) | 0.061 | 0.048 | -0.034, 0.155 | 1.259 | .208 |
|  | 2 measurements | 0.093 | 0.059 | -0.023, 0.210 | 1.567 | .117 |
|  | 3 measurements | 0.151 | 0.070 | 0.015, 0.288 | 2.169 | .030 |
|  | 1 measurement: *s* = 1 |  |  |  |  |  |
| Measurement type | Intercept (measured) | 0.081 | 0.040 | 0.002, 0.159 | 2.011 | .044 |
|  | **Rated** | **0.177** | **0.066** | **0.048, 0.306** | **2.695** | **.007** |
| Measurement type: MAT behaviors | Intercept (measured) | 0.087 | 0.041 | 0.007, 0.167 | 2.121 | .034 |
|  | **Rated** | **0.174** | **0.066** | **0.044, 0.303** | **2.630** | **.009** |
| Body masculinity type | Intercept (strength) | 0.187 | 0.031 | 0.126, 0.248 | 5.974 | <.001 |
|  | **Body shape** | **-0.099** | **0.034** | **-0.165, -0.033** | **-2.945** | **.003** |
|  | Muscle mass | -0.108 | 0.071 | -0.247, 0.031 | -1.529 | .126 |
| Body masculinity type: MAT behaviors | Intercept (strength) | 0.205 | 0.035 | 0.136, 0.274 | 5.8130 | <.001 |
|  | **Body shape** | **-0.105** | **0.040** | **-0.184, -0.026** | **-2.615** | **.009** |
|  | Muscle mass | -0.124 | 0.074 | -0.269, 0.021 | -1.676 | .094 |
| Other moderators with too few *k*/*s*: | | | | | | |
| High fertility sample: traditional vs industrialized; Non-relevant controls, Adiposity. | | | | | | |
| Reproductive domain (REP), fertility & reproductive success | | | | | | |
|  |  | B | SE | [95% CI] | *z* | *p* |
| REP measure type | Intercept (Reproductive success) | 0.170 | 0.071 | 0.032, 0.309 | 2.417 | .016 |
|  | Fertility | -0.034 | 0.081 | -0.192, 0.125 | -0.417 | .677 |
| Publication status | Intercept (published results) | 0.222 | 0.065 | 0.095, 0.349 | 3.418 | .001 |
|  | Non-published results | -0.107 | 0.075 | -0.254, 0.040 | -1.423 | .155 |
| Normality-transformed variables | Intercept (non-transformed variables) | 0.136 | 0.046 | 0.045, 0.227 | 2.928 | .003 |
|  | Transformed variables | 0.022 | 0.071 | -0.117, 0.161 | 0.311 | .756 |
| Normality-transformed variables: Fertility | Intercept (non-transformed variables) | 0.095 | 0.042 | 0.012, 0.178 | 2.246 | .025 |
|  | Transformed variables | 0.078 | 0.063 | -0.045, 0.200 | 1.239 | .215 |
| Converted effect size | Intercept (not converted) | 0.089 | 0.034 | 0.023, 0.155 | 2.635 | .008 |
|  | **Converted** | **0.143** | **0.059** | **0.028, 0.258** | **2.437** | **.015** |
| Non-relevant controls | Intercept (no non-relevant controls) | 0.136 | 0.042 | 0.054, 0.219 | 3.232 | .001 |
|  | Non-relevant controls | 0.032 | 0.082 | -0.128, 0.193 | 0.395 | .693 |
| Number of measurements | Intercept (unknown number of measurement) | 0.117 | 0.055 | 0.010, 0.224 | 2.138 | .033 |
|  | 3 measurements | 0.067 | 0.084 | -0.098, 0.232 | 0.797 | .426 |
|  | 1 measurement: *s* = 0 |  |  |  |  |  |
|  | 2 measurements: *s* = 0 |  |  |  |  |  |
| Number of measurements: Reproductive success | Intercept (unknown number of measurement) | 0.072 | 0.201 | -0.323, 0.466 | 0.356 | .722 |
|  | 3 measurements | 0.136 | 0.178 | -0.213, 0.485 | 0.763 | .446 |
|  | 1 measurement: *s* = 0 |  |  |  |  |  |
|  | 2 measurements: *s* = 0 |  |  |  |  |  |
| Body masculinity type | Intercept (strength) | 0.112 | 0.036 | 0.041, 0.183 | 3.108 | .002 |
|  | Muscle mass | 0.028 | 0.066 | -0.101, 0.158 | 0.430 | .667 |
|  | Body shape: *s* = 0 |  |  |  |  |  |
| Other moderators with too few *k*/*s*: | | | | | | |
| Sample type: low vs high fertility; Low fertility sample: students vs non-students; High fertility sample: traditional vs industrialized; Ethnicity: predominantly white vs not; Marriage system: monogamy vs non-monogamy; Peer-reviewed vs not peer-reviewed; Sexual orientation: heterosexual sample vs gay/mixed/unknown; Age controlled for; Adiposity; Measurement type: measured vs rated. | | | | | | |
| *Note.* *k* = number of observations, MAT = mating, REP = reproductive, *s* = number of samples. Moderation analyses were only run where each level of the moderator included observations from at least two studies and three independent samples. Analyses were run on the mating measures mating behaviors and mating attitudes, and the reproductive measures fertility and reproductive success when there were enough observations to do so. | | | | | | |

Supplementary File 4C

| *2D:4D: moderation analyses. The intercept shows the 'simple effect' for the reference category (specified) and the moderator effect shows the change in effect size for that category relative to the reference category. Moderators are bolded if significant after controlling for multiple comparisons, as indicated by computation of q-values. The full list of q-values can be found in Supplementary File 7.* | | | | | | | | | | | | | |
| --- | --- | --- | --- | --- | --- | --- | --- | --- | --- | --- | --- | --- | --- |
|  | Mating vs reproductive domain | | | | | | | | | | | | |
|  |  | |  | | B | | SE | [95% CI] | | | *z* | | *p* |
|  | Domain type | | Intercept (mating domain) | | 0.050 | | 0.018 | 0.015, 0.084 | | | 2.809 | | .005 |
|  |  | | Reproductive domain | | 0.007 | | 0.004 | 0.000, 0.014 | | | 1.997 | | .046 |
| Mating domain (MAT), mating behaviors & mating attitudes | | | | | | | | | | | | | |
|  | |  | | B | | SE | | | [95% CI] | *z* | | *p* | |
| MAT measure type | | Intercept (MAT behaviors) | | 0.042 | | 0.022 | | | -0.002, 0.085 | 1.870 | | .062 | |
|  | | MAT attitudes | | 0.004 | | 0.039 | | | -0.072, 0.080 | 0.097 | | .923 | |
| Low fertility sample | | Intercept (predominantly students) | | 0.036 | | 0.022 | | | -0.007, 0.078 | 1.646 | | .100 | |
|  | | Non-students/mixed/unknown | | 0.014 | | 0.042 | | | -0.069, 0.096 | 0.321 | | .749 | |
| Low fertility sample:  MAT behaviors | | Intercept (predominantly students) | | 0.033 | | 0.023 | | | -0.012, 0.077 | 1.419 | | .156 | |
|  | | Non-students/mixed/unknown | | 0.053 | | 0.049 | | | -0.042, 0.148 | 1.096 | | .273 | |
| Ethnicity | | Intercept (predominantly white) | | 0.072 | | 0.022 | | | 0.030, 0.115 | 3.333 | | .001 | |
|  | | **Mixed/other/unknown** | | **-0.080** | | **0.032** | | | **-0.143, -0.016** | **-2.462** | | **.014** | |
| Ethnicity:  MAT attitudes | | Intercept (predominantly white) | | 0.113 | | 0.063 | | | -0.010, 0.236 | 1.809 | | .071 | |
|  | | Mixed/other/unknown | | -0.128 | | 0.081 | | | -0.287, 0.032 | -1.572 | | .116 | |
| Ethnicity:  MAT behaviors | | Intercept (predominantly white) | | 0.085 | | 0.026 | | | 0.034, 0.137 | 3.245 | | .001 | |
|  | | **Mixed/other/unknown** | | **-0.088** | | **0.036** | | | **-0.158, -0.017** | **-2.423** | | **.015** | |
| Publication status | | Intercept (published results) | | 0.042 | | 0.022 | | | -0.000, 0.085 | 1.954 | | .051 | |
|  | | Non-published results | | -0.020 | | 0.040 | | | -0.098, 0.059 | -0.492 | | .623 | |
| Publication status:  MAT behaviors | | Intercept (published results) | | 0.046 | | 0.027 | | | -0.006, 0.098 | 1.719 | | .086 | |
|  | | Non-published results | | -0.017 | | 0.044 | | | -0.103, 0.068 | -0.396 | | .692 | |
| Peer-review | | Intercept (peer-reviewed) | | 0.031 | | 0.019 | | | -0.007, 0.068 | 1.615 | | .106 | |
|  | | Not peer-reviewed | | 0.034 | | 0.057 | | | -0.078, 0.146 | 0.592 | | .554 | |
| Peer-review: MAT behaviors | | Intercept (peer-reviewed) | | 0.029 | | 0.021 | | | -0.013, 0.071 | 1.349 | | .178 | |
|  | | Not peer-reviewed | | 0.070 | | 0.063 | | | -0.054, 0.193 | 1.103 | | .270 | |
| Sexual orientation | | Intercept (heterosexual sample) | | 0.014 | | 0.026 | | | -0.038, 0.065 | 0.520 | | .603 | |
|  | | Gay/mixed/unknown | | 0.035 | | 0.034 | | | -0.031, 0.102 | 1.041 | | .298 | |
| Sexual orientation: MAT behaviors | | Intercept (heterosexual sample) | | 0.024 | | 0.033 | | | -0.041, 0.089 | 0.731 | | .465 | |
|  | | Gay/mixed/unknown | | 0.024 | | 0.042 | | | -0.058, 0.105 | 0.566 | | .571 | |
| Normality-transformed variables | | Intercept (non-transformed variables) | | 0.011 | | 0.026 | | | -0.039, 0.061 | 0.443 | | .658 | |
|  | | **Transformed variables** | | **0.103** | | **0.043** | | | **0.020, 0.187** | **2.416** | | **.016** | |
| Normality-transformed variables: MAT behaviors | | Intercept (non-transformed variables) | | 0.009 | | 0.026 | | | -0.041, 0.060 | 0.368 | | .713 | |
|  | | **Transformed variables** | | **0.102** | | **0.040** | | | **0.024, 0.180** | **2.553** | | **.011** | |
| Age control | | Intercept (age controlled for) | | 0.069 | | 0.051 | | | -0.031, 0.169 | 1.358 | | .175 | |
|  | | Age not controlled for | | -0.033 | | 0.057 | | | -0.144, 0.078 | -0.583 | | .560 | |
| Age control: MAT behaviors | | Intercept (age controlled for) | | 0.077 | | 0.056 | | | -0.032, 0.186 | 1.387 | | .165 | |
|  | | Age not controlled for | | -0.025 | | 0.063 | | | -0.148, 0.098 | -0.401 | | .689 | |
| Number of measurements | | Intercept (unknown number of measurements) | | 0.021 | | 0.021 | | | -0.020, 0.061 | 1.009 | | .313 | |
|  | | 2 measurements | | -0.023 | | 0.030 | | | -0.082, 0.035 | -0.782 | | .434 | |
|  | | **3 measurements** | | **0.102** | | **0.037** | | | **0.030, 0.175** | **2.759** | | **.006** | |
|  | | 1 measurement: *s* = 1 | |  | |  | | |  |  | |  | |
| Number of measurements: MAT behaviors | | Intercept (unknown number of measurements) | | 0.021 | | 0.021 | | | -0.020, 0.061 | 1.009 | | .313 | |
|  | | 2 measurements | | -0.023 | | 0.030 | | | -0.082, 0.035 | -0.782 | | .434 | |
|  | | **3 measurements** | | **0.102** | | **0.037** | | | **0.030, 0.175** | **2.759** | | **.006** | |
|  | | 1 measurement: *s* = 1 | |  | |  | | |  |  | |  | |
| Measurement type | | Intercept (directly) | | 0.004 | | 0.032 | | | -0.058, 0.066 | 0.128 | | .898 | |
|  | | Hand scans | | 0.091 | | 0.042 | | | 0.008, 0.174 | 2.145 | | .032 | |
|  | | Unknown | | 0.005 | | 0.048 | | | -0.090, 0.098 | 0.093 | | .926 | |
|  | | Self-reported: *s* = 1 | |  | |  | | |  |  | |  | |
| Measurement type:  MAT behaviors | | Intercept (directly) | | 0.005 | | 0.032 | | | -0.059, 0.069 | 0.152 | | .879 | |
|  | | Hand scans | | 0.083 | | 0.043 | | | -0.002, 0.168 | 1.913 | | .056 | |
|  | | Unknown | | 0.000 | | 0.049 | | | -0.095, 0.096 | 0.006 | | .995 | |
|  | | Self-reported: *s* = 0 | |  | |  | | |  |  | |  | |
| Finger injuries | | Intercept (finger injuries controlled for) | | -0.002 | | 0.046 | | | -0.093, 0.088 | -0.048 | | .962 | |
|  | | Finger injuries not controlled for | | 0.046 | | 0.050 | | | -0.053, 0.144 | 0.908 | | .364 | |
| Finger injuries: MAT behaviors | | Intercept (finger injuries controlled for) | | 0.016 | | 0.047 | | | -0.077, 0.109 | 0.335 | | .738 | |
|  | | Finger injuries not controlled for | | 0.030 | | 0.053 | | | -0.074, 0.134 | 0.566 | | .572 | |
| Left vs right hand ratios | | Intercept (right 2D:4D) | | 0.039 | | 0.021 | | | -0.002, 0.080 | 1.872 | | .061 | |
|  | | Left 2D:4D | | -0.002 | | 0.005 | | | -0.013, 0.009 | -0.363 | | .717 | |
| Left vs right hand ratios: MAT attitudes | | Intercept (right 2D:4D) | | 0.006 | | 0.066 | | | -0.124, 0.136 | 0.093 | | .926 | |
|  | | Left 2D:4D | | 0.043 | | 0.060 | | | -0.076, 0.161 | 0.707 | | .479 | |
| Left vs right hand ratios: MAT behaviors | | Intercept (right 2D:4D) | | 0.037 | | 0.029 | | | -0.021, 0.094 | 1.253 | | .210 | |
|  | | Left 2D:4D | | 0.014 | | 0.032 | | | -0.048, 0.077 | 0.449 | | .653 | |
| Other moderators with too few *k*/*s*: | | | | | | | | | | | | | |
| Sample type: low vs high fertility; High fertility sample: traditional vs industrialized; Marriage system: monogamy vs non-monogamy; Converted effect size; Non-relevant controls. | | | | | | | | | | | | | |
| Reproductive domain (REP), fertility & reproductive success | | | | | | | | | | | | | |
|  | |  | | B | | SE | | | [95% CI] | *z* | | *p* | |
| REP measure type | | Intercept (reproductive success) | | 0.170 | | 0.066 | | | 0.042, 0.301 | 2.592 | | .010 | |
|  | | Fertility | | -0.135 | | 0.078 | | | -0.289, 0.016 | -1.751 | | .080 | |
| Sample type | | Intercept (low fertility) | | 0.075 | | 0.057 | | | -0.037, 0.187 | 1.319 | | .187 | |
|  | | High fertility | | 0.002 | | 0.072 | | | -0.143, 0.140 | -0.022 | | .983 | |
| High fertility sample | | Intercept (traditional) | | 0.028 | | 0.089 | | | -0.145, 0.202 | 0.320 | | .749 | |
|  | | Industrialized | | 0.110 | | 0.126 | | | -0.137, 0.357 | 0.874 | | .382 | |
| Ethnicity | | Intercept (predominantly white) | | 0.066 | | 0.055 | | | -0.042, 0.175 | 1.196 | | .232 | |
|  | | Mixed/other/unknown | | 0.016 | | 0.072 | | | -0.124, 0.157 | 0.227 | | .821 | |
| Marriage system | | Intercept (monogamy) | | 0.054 | | 0.042 | | | -0.027, 0.136 | 1.304 | | .192 | |
|  | | Non-monogamy | | 0.116 | | 0.089 | | | -0.058, 0.290 | 1.306 | | .192 | |
| Publication status | | Intercept (published results) | | 0.105 | | 0.040 | | | 0.026, 0.184 | 2.601 | | .009 | |
|  | | Non-published results | | -0.166 | | 0.093 | | | -0.348, 0.015 | -1.795 | | .073 | |
| Sexual orientation | | Intercept (heterosexual sample) | | 0.039 | | 0.066 | | | -0.091, 0.169 | 0.584 | | .560 | |
|  | | Gay/mixed/unknown | | 0.062 | | 0.087 | | | -0.109, 0.233 | 0.707 | | .480 | |
| Sexual orientation: Fertility | | Intercept (heterosexual sample) | | 0.037 | | 0.079 | | | -0.119, 0.192 | 0.462 | | .644 | |
|  | | Gay/mixed/unknown | | -0.019 | | 0.125 | | | -0.264, 0.225 | -0.155 | | .877 | |
| Normality-transformed variables | | Intercept (non-transformed variables) | | 0.031 | | 0.036 | | | -0.040, 0.101 | 0.851 | | .395 | |
|  | | Transformed variables | | 0.138 | | 0.067 | | | 0.006, 0.269 | 2.054 | | .040 | |
| Age control | | Intercept (age controlled for) | | 0.070 | | 0.037 | | | -0.002, 0.142 | 1.915 | | .056 | |
|  | | Age not controlled for | | 0.077 | | 0.085 | | | -0.088, 0.243 | 0.916 | | .360 | |
| Measurement type | | Intercept (directly) | | 0.072 | | 0.064 | | | -0.053, 0.197 | 1.133 | | .257 | |
|  | | Hand scans | | 0.053 | | 0.106 | | | -0.155, 0.261 | 0.502 | | .616 | |
|  | | Self-reported: *s* = 1 | |  | |  | | |  |  | |  | |
|  | | Unknown: *s* = 0 | |  | |  | | |  |  | |  | |
| Finger injuries | | Intercept (finger injuries controlled for) | | 0.136 | | 0.062 | | | 0.014, 0.257 | 2.190 | | .029 | |
|  | | Finger injuries not controlled for | | -0.104 | | 0.081 | | | -0.262, -0.054 | -1.292 | | .196 | |
| Left vs right hand ratios | | Intercept (right 2D:4D) | | 0.070 | | 0.044 | | | -0.017, 0.156 | 1.584 | | .113 | |
|  | | Left 2D:4D | | 0.004 | | 0.004 | | | -0.005, 0.012 | 0.840 | | .401 | |
| Left vs right hand ratios: Fertility | | Intercept (right 2D:4D) | | 0.031 | | 0.050 | | | -0.067, 0.128 | 0.618 | | .537 | |
|  | | Left 2D:4D | | 0.004 | | 0.004 | | | -0.005, 0.012 | 0.853 | | .394 | |
| Other moderators with too few *k*/*s*: | | | | | | | | | | | | | |
| Low fertility sample: students vs non-students; Peer-reviewed vs not peer-reviewed; Converted effect size; Non-relevant controls; Number of measurements; Left vs right hand. | | | | | | | | | | | | | |
| *Note.* *k* = number of observations, MAT = mating, REP = reproductive, *s* = number of samples. Moderation analyses were only run where each level of the moderator included observations from at least two studies and three independent samples. Analyses were run on the mating measures mating behaviors and mating attitudes, and the reproductive measures fertility and reproductive success when there were enough observations to do so. | | | | | | | | | | | | | |

Supplementary File 4D

| *Voice pitch: moderation analyses. The intercept shows the 'simple effect' for the reference category (specified) and the moderator effect shows the change in effect size for that category relative to the reference category. Moderators are bolded if significant after controlling for multiple comparisons, as indicated by computation of q-values. The full list of q-values can be found in Supplementary File 7.* | | | | | | | |
| --- | --- | --- | --- | --- | --- | --- | --- |
|  | Mating vs reproductive domain | | | | | | |
|  |  |  | B | SE | [95% CI] | *z* | *p* |
|  | Domain type | Intercept (mating domain) | 0.132 | 0.037 | 0.061, 0.204 | 3.610 | <.001 |
|  |  | Reproductive domain | 0.004 | 0.075 | -0.143, 0.151 | 0.059 | .953 |
| Mating domain (MAT), mating behaviors & mating attitudes | | | | | | | |
| Moderators with too few *k*/*s* (i.e. all potential moderators): | | | | | | | |
| MAT measure type: behaviors vs attitudes; Sample type: low vs high fertility; Low fertility sample: predominantly students vs non-students; High fertility sample: traditional vs industrialized; Ethnicity: predominantly white vs not; Marriage system: monogamy vs non-monogamy; Publication status: published results; Peer-reviewed; Sexual orientation: heterosexual sample vs gay/mixed/unknown; Normality-transformed variables; Converted effect size; Age controlled for; Non-relevant controls; Sex of experimenter; Illness; Smoker; Condition: baseline vs competition vs courtship. | | | | | | | |
| Reproductive domain (REP), fertility & reproductive success | | | | | | | |
| Moderators with too few *k*/*s* (i.e. all potential moderators): | | | | | | | |
| REP measure type: reproductive success vs fertility; Sample type: low vs high fertility; Low fertility sample: predominantly students vs non-students; High fertility sample: traditional vs industrialized; Ethnicity: predominantly white vs not; Marriage system: monogamy vs non-monogamy; Publication status: published results; Peer-reviewed; Sexual orientation: heterosexual sample vs gay/mixed/unknown; Normality-transformed variables; Converted effect size; Age controlled for; Non-relevant controls; Sex of experimenter; Illness; Smoker; Condition: baseline vs competition vs courtship. | | | | | | | |
| *Note.* *k* = number of observations, MAT = mating, REP = reproductive, *s* = number of samples. Moderation analyses were only run where each level of the moderator included observations from at least two studies and three independent samples. Analyses were run on the mating measures mating behaviors and mating attitudes, and the reproductive measures fertility and reproductive success when there were enough observations to do so. | | | | | | | |

Supplementary File 4E

| *Height: moderation analyses. The intercept shows the 'simple effect' for the reference category (specified) and the moderator effect shows the change in effect size for that category relative to the reference category. Moderators are bolded if significant after controlling for multiple comparisons, as indicated by computation of q-values. The full list of q-values can be found in Supplementary File 7.* | | | | | | | | | | | | | |
| --- | --- | --- | --- | --- | --- | --- | --- | --- | --- | --- | --- | --- | --- |
|  | Mating vs reproductive domain | | | | | | | | | | | | |
|  |  | |  | | B | | SE | [95% CI] | | | *z* | | *p* |
|  | Domain type | | Intercept (mating domain) | | 0.049 | | 0.020 | 0.010, 0.088 | | | 2.475 | | .013 |
|  |  | | Reproductive domain | | -0.032 | | 0.017 | -0.066, 0.001 | | | -1.900 | | .057 |
| Mating domain (MAT), mating behaviors & mating attitudes | | | | | | | | | | | | | |
|  | |  | | B | | SE | | | [95% CI] | *z* | | *p* | |
| MAT measure type | | Intercept (MAT behaviors) | | 0.054 | | 0.015 | | | 0.024, 0.084 | 3.504 | | .001 | |
|  | | MAT attitudes | | 0.000 | | 0.021 | | | -0.041, 0.041 | 0.007 | | .995 | |
| Sample type | | Intercept (low fertility) | | 0.055 | | 0.016 | | | 0.024, 0.086 | 3.456 | | .001 | |
|  | | High fertility | | 0.031 | | 0.062 | | | -0.092, 0.153 | 0.492 | | .623 | |
| Sample type: MAT behaviors | | Intercept (low fertility) | | 0.051 | | 0.018 | | | 0.017, 0.086 | 2.923 | | .004 | |
|  | | High fertility | | 0.034 | | 0.063 | | | -0.090, 0.158 | 0.541 | | .588 | |
| Low fertility sample | | Intercept (predominantly students) | | 0.065 | | 0.024 | | | 0.018, 0.111 | 2.722 | | .007 | |
|  | | Non-students/mixed/unknown | | -0.018 | | 0.033 | | | -0.082, 0.047 | -0.537 | | .591 | |
| Low fertility sample:  MAT attitudes | | Intercept (predominantly students) | | 0.045 | | 0.053 | | | -0.060, 0.150 | 0.843 | | .399 | |
|  | | Non-students/mixed/unknown | | -0.020 | | 0.058 | | | -0.135, 0.094 | -0.346 | | .729 | |
| Low fertility sample:  MAT behaviors | | Intercept (predominantly students) | | 0.057 | | 0.026 | | | 0.006, 0.109 | 2.171 | | .030 | |
|  | | Non-students/mixed/unknown | | -0.010 | | 0.037 | | | -0.082, 0.062 | -0.284 | | .777 | |
| Ethnicity | | Intercept (predominantly white) | | 0.073 | | 0.023 | | | 0.028, 0.118 | 3.153 | | .002 | |
|  | | Mixed/other/unknown | | -0.029 | | 0.031 | | | -0.090, 0.032 | -0.918 | | .359 | |
| Ethnicity: MAT behaviors | | Intercept (predominantly white) | | 0.072 | | 0.026 | | | 0.021, 0.123 | 2.761 | | .006 | |
|  | | Mixed/other/unknown | | -0.031 | | 0.035 | | | -0.098, 0.037 | -0.890 | | .374 | |
| Publication status | | Intercept (published results) | | 0.067 | | 0.030 | | | 0.009, 0.126 | 2.269 | | .023 | |
|  | | Non-published results | | -0.014 | | 0.035 | | | -0.082, 0.054 | -0.403 | | .687 | |
| Publication status:  MAT behaviors | | Intercept (published results) | | 0.074 | | 0.030 | | | 0.016, 0.133 | 2.486 | | .013 | |
|  | | Non-published results | | -0.030 | | 0.036 | | | -0.100, 0.040 | -0.833 | | .405 | |
| Peer-review | | Intercept (peer-reviewed) | | 0.055 | | 0.016 | | | 0.023, 0.087 | 3.348 | | .001 | |
|  | | Not peer-reviewed | | 0.031 | | 0.058 | | | -0.083, 0.145 | 0.534 | | .593 | |
| Peer-review: MAT behaviors | | Intercept (peer-reviewed) | | 0.050 | | 0.018 | | | 0.016, 0.085 | 2.867 | | .004 | |
|  | | Not peer-reviewed | | 0.052 | | 0.066 | | | -0.076, 0.181 | 0.799 | | .424 | |
| Sexual orientation | | Intercept (heterosexual sample) | | 0.041 | | 0.021 | | | -0.000, 0.082 | 1.954 | | .051 | |
|  | | Gay/mixed/unknown | | 0.038 | | 0.032 | | | -0.025, 0.101 | 1.180 | | .238 | |
| Sexual orientation: MAT behaviors | | Intercept (heterosexual sample) | | 0.035 | | 0.023 | | | -0.011, 0.081 | 1.507 | | .132 | |
|  | | Gay/mixed/unknown | | 0.042 | | 0.035 | | | -0.027, 0.111 | 1.199 | | .231 | |
| Normality-transformed variables | | Intercept (non-transformed variables) | | 0.049 | | 0.020 | | | 0.011, 0.087 | 2.521 | | .012 | |
|  | | Transformed variables | | 0.031 | | 0.031 | | | -0.030, 0.091 | 0.995 | | .320 | |
| Normality-transformed variables: MAT behaviors | | Intercept (non-transformed variables) | | 0.039 | | 0.022 | | | -0.004, 0.081 | 1.796 | | .073 | |
|  | | Transformed variables | | 0.053 | | 0.036 | | | -0.017, 0.122 | 1.480 | | .139 | |
| Converted effect size | | Intercept (not converted) | | 0.052 | | 0.017 | | | 0.019, 0.086 | 3.068 | | .002 | |
|  | | Converted | | 0.048 | | 0.043 | | | -0.036, 0.133 | 1.118 | | .264 | |
| Converted effect size: MAT behaviors | | Intercept (not converted) | | 0.046 | | 0.019 | | | 0.010, 0.082 | 2.481 | | .013 | |
|  | | Converted | | 0.055 | | 0.044 | | | -0.031, 0.141 | 1.244 | | .214 | |
| Age control | | Intercept (age controlled for) | | 0.056 | | 0.018 | | | 0.021, 0.090 | 3.160 | | .002 | |
|  | | Age not controlled for | | 0.013 | | 0.018 | | | -0.022, 0.048 | 0.708 | | .479 | |
| Age control: MAT behaviors | | Intercept (age controlled for) | | 0.051 | | 0.017 | | | 0.018, 0.084 | 2.993 | | .003 | |
|  | | Age not controlled for | | 0.012 | | 0.018 | | | -0.023, 0.047 | 0.663 | | .507 | |
| Number of measurements | | Intercept (unknown number of measurements) | | 0.047 | | 0.021 | | | 0.006, 0.088 | 2.255 | | .024 | |
|  | | 2 measurements | | 0.096 | | 0.057 | | | -0.015, 0.207 | 1.702 | | .089 | |
|  | | 1 measurement: *s* = 1 | |  | |  | | |  |  | |  | |
| Number of measurements: MAT behaviors | | Intercept (unknown number of measurements) | | 0.051 | | 0.023 | | | 0.006, 0.095 | 2.231 | | .026 | |
|  | | 2 measurements | | 0.088 | | 0.070 | | | -0.049, 0.226 | 1.264 | | .206 | |
|  | | 1 measurement: *s* = 1 | |  | |  | | |  |  | |  | |
| Measurement type | | Intercept (measured) | | 0.057 | | 0.021 | | | 0.016, 0.098 | 2.705 | | .007 | |
|  | | Self-reported | | 0.001 | | 0.036 | | | -0.069, 0.071 | 0.035 | | .972 | |
|  | | Unknown measurement type | | 0.010 | | 0.067 | | | -0.122, 0.142 | 0.144 | | .886 | |
| Measurement type:  MAT behaviors | | Intercept (measured) | | 0.054 | | 0.022 | | | 0.011, 0.097 | 2.455 | | .014 | |
|  | | Self-reported | | -0.003 | | 0.040 | | | -0.080, 0.075 | -0.073 | | .942 | |
|  | | Unknown measurement type | | 0.026 | | 0.079 | | | -0.128, 0.180 | 0.329 | | .742 | |
| Other moderators with too few *k*/*s*: | | | | | | | | | | | | | |
| High fertility sample: traditional vs industrialized; Marriage system: monogamy vs non-monogamy; Non-relevant controls. | | | | | | | | | | | | | |
| Reproductive domain (REP), fertility & reproductive success | | | | | | | | | | | | | |
|  | |  | | B | | SE | | | [95% CI] | *z* | | *p* | |
| REP measure type | | Intercept (reproductive success) | | -0.031 | | 0.053 | | | -0.135, 0.073 | -0.584 | | .559 | |
|  | | Fertility | | 0.045 | | 0.054 | | | -0.061, 0.150 | 0.831 | | .406 | |
| Sample type | | Intercept (low fertility) | | -0.037 | | 0.044 | | | -0.123, 0.050 | -0.825 | | .409 | |
|  | | High fertility | | 0.071 | | 0.057 | | | -0.041, 0.182 | 1.243 | | .214 | |
| Sample type: Fertility | | Intercept (low fertility) | | -0.037 | | 0.035 | | | -0.105, 0.031 | -1.060 | | .289 | |
|  | | High fertility | | 0.090 | | 0.048 | | | -0.004, 0.185 | 1.878 | | .060 | |
| High fertility sample | | Intercept (traditional) | | 0.029 | | 0.051 | | | -0.071, 0.130 | 0.575 | | .565 | |
|  | | Industrialized | | 0.011 | | 0.081 | | | -0.147, 0.170 | 0.141 | | .888 | |
| High fertility sample: Fertility | | Intercept (traditional) | | 0.048 | | 0.042 | | | -0.035, 0.131 | 1.132 | | .258 | |
|  | | Industrialized | | 0.016 | | 0.057 | | | -0.096, 0.129 | 0.287 | | .774 | |
| Ethnicity | | Intercept (predominantly white) | | 0.038 | | 0.047 | | | -0.054, 0.130 | 0.808 | | .419 | |
|  | | Mixed/other/unknown | | -0.049 | | 0.059 | | | -0.165, 0.066 | -0.839 | | .401 | |
| Ethnicity: Fertility | | Intercept (predominantly white) | | 0.037 | | 0.040 | | | -0.042, 0.116 | 0.924 | | .355 | |
|  | | Mixed/other/unknown | | -0.044 | | 0.053 | | | -0.147, 0.059 | -0.838 | | .402 | |
| Marriage system | | Intercept (monogamy) | | -0.010 | | 0.034 | | | -0.078, 0.057 | -0.304 | | .761 | |
|  | | Non-monogamy | | 0.052 | | 0.060 | | | -0.066, 0.169 | 0.862 | | .389 | |
| Marriage system: Fertility | | Intercept (monogamy) | | -0.001 | | 0.030 | | | -0.059, 0.057 | -0.034 | | .973 | |
|  | | Non-monogamy | | 0.052 | | 0.061 | | | -0.067, 0.171 | 0.860 | | .390 | |
| Publication status | | Intercept (published results) | | -0.023 | | 0.038 | | | -0.098, 0.052 | -0.601 | | .548 | |
|  | | Non-published results | | 0.064 | | 0.056 | | | -0.046, 0.175 | 1.139 | | .255 | |
| Publication status: Fertility | | Intercept (published results) | | -0.018 | | 0.036 | | | -0.088, 0.052 | -0.501 | | .617 | |
|  | | Non-published results | | 0.063 | | 0.052 | | | -0.040, 0.165 | 1.202 | | .229 | |
| Sexual orientation | | Intercept (heterosexual sample) | | -0.092 | | 0.048 | | | -0.185, 0.001 | -1.939 | | .053 | |
|  | | **Gay/mixed/unknown** | | **0.135** | | **0.056** | | | **0.026, 0.245** | **2.417** | | **.016** | |
| Sexual orientation: Fertility | | Intercept (heterosexual sample) | | -0.070 | | 0.041 | | | -0.151, 0.011 | -1.702 | | .089 | |
|  | | Gay/mixed/unknown | | 0.117 | | 0.050 | | | 0.019, 0.214 | 2.342 | | .019 | |
| Normality-transformed variables | | Intercept (non-transformed variables) | | -0.005 | | 0.033 | | | -0.069, 0.060 | -0.149 | | .881 | |
|  | | Transformed variables | | 0.049 | | 0.065 | | | -0.078, 0.175 | 0.756 | | .450 | |
| Normality-transformed variables: Fertility | | Intercept (non-transformed variables) | | -0.002 | | 0.030 | | | -0.060, 0.056 | -0.081 | | .936 | |
|  | | Transformed variables | | 0.067 | | 0.064 | | | -0.059, 0.193 | 1.036 | | .300 | |
| Converted effect size | | Intercept (not converted) | | 0.022 | | 0.034 | | | -0.045, 0.089 | 0.645 | | .519 | |
|  | | Converted | | -0.048 | | 0.058 | | | -0.161, 0.065 | -0.833 | | .405 | |
| Converted effect size: Fertility | | Intercept (not converted) | | 0.033 | | 0.032 | | | -0.029, 0.095 | 1.051 | | .293 | |
|  | | Converted | | -0.072 | | 0.058 | | | -0.186, 0.041 | -1.246 | | .213 | |
| Converted effect size: Reproductive success | | Intercept (not converted) | | -0.005 | | 0.132 | | | -0.264, 0.254 | -0.037 | | .971 | |
|  | | Converted | | -0.086 | | 0.187 | | | -0.452, 0.280 | -0.463 | | .644 | |
| Age control | | Intercept (age controlled for) | | 0.005 | | 0.034 | | | -0.061, 0.071 | 0.154 | | .878 | |
|  | | Age not controlled for | | -0.032 | | 0.083 | | | -0.195, 0.131 | -0.386 | | .699 | |
| Non-relevant controls | | Intercept (no non-relevant controls) | | 0.012 | | 0.032 | | | -0.050, 0.075 | 0.387 | | .699 | |
|  | | Non-relevant controls | | -0.029 | | 0.068 | | | -0.161, 0.103 | -0.428 | | .668 | |
| Non-relevant controls: Fertility | | Intercept (no non-relevant controls) | | 0.017 | | 0.030 | | | -0.041, 0.076 | 0.585 | | .559 | |
|  | | Non-relevant controls | | -0.033 | | 0.073 | | | -0.176, 0.111 | -0.443 | | .657 | |
| Non-relevant controls: Reproductive success | | Intercept (no non-relevant controls) | | 0.017 | | 0.116 | | | -0.210, 0.244 | 0.150 | | .881 | |
|  | | Non-relevant controls | | -0.166 | | 0.184 | | | -0.526, 0.194 | -0.902 | | .367 | |
| Measurement type | | Intercept (measured) | | -0.007 | | 0.039 | | | -0.083, 0.070 | -0.167 | | .867 | |
|  | | Self-reported | | 0.012 | | 0.065 | | | -0.116, 0.140 | 0.178 | | .859 | |
|  | | Unknown: *s* = 2 | |  | |  | | |  |  | |  | |
| Measurement type:  Fertility | | Intercept (measured) | | -0.001 | | 0.038 | | | -0.076, 0.073 | -0.037 | | .971 | |
|  | | Self-reported | | 0.006 | | 0.058 | | | -0.108, 0.121 | 0.105 | | .916 | |
|  | | Unknown: *s* = 2 | |  | |  | | |  |  | |  | |
| Other moderators with too few *k*/*s*: | | | | | | | | | | | | | |
| Low fertility sample: predominantly students vs non-students; Peer-reviewed; Number of measurements. | | | | | | | | | | | | | |
| *Note.* *k* = number of observations, MAT = mating, REP = reproductive, *s* = number of samples. Moderation analyses were only run where each level of the moderator included observations from at least two studies and three independent samples. Analyses were run on the mating measures mating behaviors and mating attitudes, and the reproductive measures fertility and reproductive success when there were enough observations to do so. | | | | | | | | | | | | | |

Supplementary File 4F

| *Testosterone levels: moderation analyses. The intercept shows the 'simple effect' for the reference category (specified) and the moderator effect shows the change in effect size for that category relative to the reference category. Moderators are bolded if significant after controlling for multiple comparisons, as indicated by computation of q-values. The full list of q-values can be found in Supplementary File 7.* | | | | | | | | | | | | | | | | |
| --- | --- | --- | --- | --- | --- | --- | --- | --- | --- | --- | --- | --- | --- | --- | --- | --- |
|  | Mating vs reproductive domain | | | | | | | | | | | | | | | |
|  |  | |  | | B | | SE | | [95% CI] | | | *z* | | | *p* | |
|  | Domain type | | Intercept (Mating domain) | | 0.093 | | 0.014 | | 0.067, 0.120 | | | 6.893 | | | <.001 | |
|  |  | | Reproductive domain | | -0.063 | | 0.060 | | -0.181, 0.054 | | | -1.059 | | | .290 | |
| Mating domain (MAT), mating behaviors & mating attitudes | | | | | | | | | | | | | | | | |
|  | |  | | B | | SE | | | | [95% CI] | *z* | | | *p* | | |
| MAT measure type | | Intercept (MAT behaviors) | | 0.087 | | 0.016 | | 0.056, 0.118 | | | | | 5.460 | | | <.001 |
|  | | MAT attitudes | | 0.015 | | 0.027 | | -0.038, 0.068 | | | | | 0.569 | | | .569 |
| Low fertility sample | | Intercept (predominantly students) | | 0.096 | | 0.020 | | 0.057, 0.136 | | | | | 4.755 | | | <.001 |
|  | | Non-students/mixed/unknown | | 0.012 | | 0.041 | | -0.069, 0.094 | | | | | 0.298 | | | .766 |
| Low fertility sample:  MAT behaviors | | Intercept (predominantly students) | | 0.080 | | 0.024 | | 0.032, 0.128 | | | | | 3.288 | | | .001 |
|  | | Non-students/mixed/unknown | | 0.024 | | 0.039 | | -0.053, 0.101 | | | | | 0.610 | | | .542 |
| Ethnicity | | Intercept (predominantly white) | | 0.104 | | 0.023 | | 0.058, 0.150 | | | | | 4.436 | | | <.001 |
|  | | Mixed/other/unknown | | -0.016 | | 0.030 | | -0.074, 0.043 | | | | | -0.530 | | | .596 |
| Ethnicity: MAT attitudes | | Intercept (predominantly white) | | 0.126 | | 0.060 | | 0.008, 0.245 | | | | | 2.090 | | | .037 |
|  | | Mixed/other/unknown | | -0.045 | | 0.074 | | -0.189, 0.099 | | | | | -0.614 | | | .540 |
| Ethnicity: MAT behaviors | | Intercept (predominantly white) | | 0.099 | | 0.022 | | 0.056, 0.142 | | | | | 4.497 | | | <.001 |
|  | | Mixed/other/unknown | | -0.024 | | 0.025 | | -0.074, 0.026 | | | | | -0.943 | | | .346 |
| Publication status | | Intercept (published results) | | 0.097 | | 0.017 | | 0.064, 0.130 | | | | | 5.736 | | | <.001 |
|  | | Non-published results | | 0.012 | | 0.039 | | -0.088, 0.065 | | | | | 0.303 | | | .762 |
| Publication status:  MAT behaviors | | Intercept (published results) | | 0.082 | | 0.014 | | 0.054, 0.110 | | | | | 5.703 | | | <.001 |
|  | | Non-published results | | 0.019 | | 0.041 | | -0.061, 0.098 | | | | | 0.465 | | | .642 |
| Sexual orientation | | Intercept (heterosexual sample) | | 0.125 | | 0.016 | | 0.092, 0.157 | | | | | 7.578 | | | <.001 |
|  | | **Gay/mixed/unknown** | | **-0.059** | | **0.020** | | **-0.098, -0.021** | | | | | **-2.994** | | | **.003** |
| Sexual orientation: MAT attitudes | | Intercept (heterosexual sample) | | 0.165 | | 0.059 | | 0.050, 0.281 | | | | | 2.803 | | | .005 |
|  | | Gay/mixed/unknown | | -0.108 | | 0.076 | | -0.256, 0.041 | | | | | -1.419 | | | .156 |
| Sexual orientation: MAT behaviors | | Intercept (heterosexual sample) | | 0.110 | | 0.020 | | 0.071, 0.149 | | | | | 5.529 | | | <.001 |
|  | | Gay/mixed/unknown | | -0.042 | | 0.024 | | -0.089, 0.005 | | | | | -1.751 | | | .080 |
| Normality-transformed variables | | Intercept (non-transformed variables) | | 0.073 | | 0.010 | | 0.053, 0.093 | | | | | 7.172 | | | <.001 |
|  | | **Transformed variables** | | **0.057** | | **0.023** | | **0.011, 0.103** | | | | | **2.445** | | | **.015** |
| Normality-transformed variables: MAT behaviors | | Intercept (non-transformed variables) | | 0.074 | | 0.012 | | 0.050, 0.098 | | | | | 5.995 | | | <.001 |
|  | | Transformed variables | | 0.036 | | 0.028 | | -0.018, 0.091 | | | | | 1.306 | | | .192 |
| Converted effect size | | Intercept (not converted) | | 0.091 | | 0.016 | | 0.061, 0.121 | | | | | 5.863 | | | <.001 |
|  | | Converted | | 0.029 | | 0.043 | | -0.056, 0.114 | | | | | 0.665 | | | .506 |
| Converted effect size: MAT behaviors | | Intercept (not converted) | | 0.078 | | 0.013 | | 0.054, 0.103 | | | | | 6.227 | | | <.001 |
|  | | Converted | | 0.058 | | 0.046 | | -0.032, 0.147 | | | | | 1.266 | | | .206 |
| Age control | | Intercept (age controlled for) | | 0.098 | | 0.022 | | 0.054, 0.142 | | | | | 4.393 | | | <.001 |
|  | | Age not controlled for | | -0.011 | | 0.032 | | -0.074, 0.052 | | | | | -0.346 | | | .730 |
| Age control: MAT behaviors | | Intercept (age controlled for) | | 0.107 | | 0.029 | | 0.050, 0.163 | | | | | 3.690 | | | <.001 |
|  | | Age not controlled for | | -0.029 | | 0.041 | | -0.109, 0.051 | | | | | -0.708 | | | .479 |
| Non-relevant controls | | Intercept (no non-relevant controls) | | 0.088 | | 0.015 | | 0.059, 0.117 | | | | | 5.954 | | | <.001 |
|  | | Non-relevant controls | | 0.042 | | 0.041 | | -0.039, 0.122 | | | | | 1.012 | | | .312 |
| Time of day | | Intercept (AM) | | 0.090 | | 0.023 | | 0.045, 0.135 | | | | | 3.899 | | | <.001 |
|  | | PM | | 0.016 | | 0.028 | | -0.038, 0.070 | | | | | 0.568 | | | .570 |
|  | | Mixed/unknown | | -0.014 | | 0.046 | | -0.105, 0.076 | | | | | -0.309 | | | .757 |
| Time of day: MAT attitudes | | Intercept (AM) | | 0.069 | | 0.045 | | -0.019, 0.157 | | | | | 1.543 | | | .123 |
|  | | PM | | 0.038 | | 0.047 | | -0.055, 0.130 | | | | | 0.794 | | | .427 |
|  | | Mixed/unknown: *s* = 2 | |  | |  | |  | | | | |  | | |  |
| Time of day: MAT behaviors | | Intercept (AM) | | 0.088 | | 0.024 | | 0.041, 0.134 | | | | | 3.672 | | | <.001 |
|  | | PM | | 0.003 | | 0.036 | | -0.068, 0.074 | | | | | 0.087 | | | .931 |
|  | | Mixed/unknown | | -0.017 | | 0.049 | | -0.114, 0.079 | | | | | -0.350 | | | .726 |
| Blood contamination | | Intercept (checked) | | 0.142 | | 0.031 | | 0.080, 0.203 | | | | | 4.533 | | | <.001 |
|  | | Not checked/unknown | | -0.061 | | 0.036 | | -0.133, 0.010 | | | | | -1.694 | | | .090 |
| Blood contamination: MAT behaviors | | Intercept (checked) | | 0.130 | | 0.032 | | 0.067, 0.193 | | | | | 4.036 | | | <.001 |
|  | | Not checked/unknown | | -0.064 | | 0.040 | | -0.141, 0.014 | | | | | -1.599 | | | .110 |
| Fatherhood status | | Intercept (non-fathers) | | 0.069 | | 0.025 | | 0.021, 0.117 | | | | | 2.831 | | | .005 |
|  | | Mixed/unknown | | 0.040 | | 0.031 | | -0.021, 0.101 | | | | | 1.286 | | | .198 |
|  | | Fathers: *s* = 0 | |  | |  | |  | | | | |  | | |  |
| Fatherhood status: MAT attitudes | | Intercept (non-fathers) | | 0.072 | | 0.054 | | -0.033, 0.177 | | | | | 1.339 | | | .181 |
|  | | Mixed/unknown | | 0.061 | | 0.081 | | -0.097, 0.219 | | | | | 0.755 | | | .450 |
|  | | Fathers: *s* = 0 | |  | |  | |  | | | | |  | | |  |
| Fatherhood status: MAT behaviors | | Intercept (non-fathers) | | 0.055 | | 0.031 | | -0.005, 0.115 | | | | | 1.792 | | | .073 |
|  | | Mixed/unknown | | 0.043 | | 0.036 | | -0.027, 0.114 | | | | | 1.203 | | | .229 |
|  | | Fathers: *s* = 0 | |  | |  | |  | | | | |  | | |  |
| Other moderators with too few *k*/*s*: | | | | | | | | | | | | | | | | |
| Sample type: low vs high fertility; High fertility sample: traditional vs industrialized; Marriage system: monogamy vs non-monogamy; Peer-reviewed; How assayed: blood vs saliva; Relationship status | | | | | | | | | | | | | | | | |
| Reproductive domain (REP), fertility & reproductive success | | | | | | | | | | | | | | | | |
| Moderators with too few *k*/*s* (i.e. all potential moderators): | | | | | | | | | | | | | | | | |
| REP measure type (reproductive success vs fertility; Sample type: low vs high fertility; Low fertility sample: predominantly students vs non-students; High fertility sample: traditional vs industrialized; Ethnicity: predominantly white vs not; Marriage system: monogamy vs non-monogamy; Publication status: published results; Peer-reviewed; Sexual orientation: heterosexual sample vs gay/mixed/unknown; Normality-transformed variables; Converted effect size; Age controlled for; Non-relevant controls; How assayed: blood vs saliva; Time of day: AM vs PM vs mixed/unknown; Fatherhood status; Relationship status | | | | | | | | | | | | | | | | |
| *Note.* *k* = number of observations, MAT = mating, REP = reproductive, *s* = number of samples. Moderation analyses were only run where each level of the moderator included observations from at least two studies and three independent samples. Analyses were run on the mating measures mating behaviors and mating attitudes, and the reproductive measures fertility and reproductive success when there were enough observations to do so. | | | | | | | | | | | | | | | | |

Supplementary File 5A

| *Mating domain, reproductive domain, and offspring mortality domain predicted by global masculinity. Pearson’s r (95% CI); p value for meta-analytic effect, q-value (correcting for multiple comparisons); number of observations (k), samples (s), and unique participants (n); test for heterogeneity (Q), p value for heterogeneity. Statistically significant meta-analytic associations are bolded if still significant after controlling for multiple comparisons.* | | | |
| --- | --- | --- | --- |
|  | Mating domain | Reproductive domain | Offspring mortality domain |
| Sample |  |  |  |
| All samples | ***r* = .090 (0.071, 0.110), *p* < .001, *q* = .001** *k* = 371, *s* = 70, *n* = 117481 Q(df = 370) = 1108.213, *p* < .001 | *r* = .047 (0.004, 0.090),  *p* = .033, *q* = .080 *k* = 81, *s* = 36, *n* = 107848 Q(df = 80) = 628.883, *p* < .001 | *r* = .002 (-0.011, 0.015), *p* = .782, *q* = .475 *k* = 22, *s* = 13, *n* = 21991 Q(df = 21) = 14.765, *p* = .835 |
| *Note.* k = number of observations; n = number of unique participants; Q = Cochran’s Q test of heterogeneity; s = number of samples. | | | |

Supplementary File 5B

| *Global masculinity: moderation analyses. The intercept shows the 'simple effect' for the reference category (specified) and the moderator effect shows the change in effect size for that category relative to the reference category. Moderators are bolded if significant after controlling for multiple comparisons, as indicated by computation of q-values. The full list of q-values can be found in Supplementary File 7.* | | | | | | | |
| --- | --- | --- | --- | --- | --- | --- | --- |
|  | Global masculinity | | | | | | |
|  |  |  | B | SE | [95% CI] | *z* | *p* |
|  | Domain type | **Intercept (mating domain)** | **0.083** | **0.010** | **0.062, 0.103** | **8.022** | **<.001** |
|  |  | Reproductive domain | 0.004 | 0.003 | -0.003, 0.011 | 1.209 | .227 |
|  |  | Offspring mortality domain | -0.054 | 0.030 | -0.113, 0.005 | -1.794 | .073 |
| Mating domain: moderation analyses of type of masculinity | | | | | | | |
|  |  |  | B | SE | [95% CI] | *z* | *p* |
|  | Masculinity type | **Intercept (body masculinity)** | **0.108** | **0.012** | **0.085, 0.132** | **8.951** | **<.001** |
|  |  | **2D:4D** | **-0.060** | **0.018** | **-0.096, -0.025** | **-3.325** | **.001** |
|  |  | **Facial masculinity** | **-0.054** | **0.023** | **-0.099, -0.009** | **-2.354** | **.019** |
|  |  | **Height** | **-0.020** | **0.009** | **-0.037, -0.002** | **-2.235** | **.025** |
|  |  | Testosterone levels | -0.009 | 0.020 | -0.047, 0.030 | -0.452 | .652 |
|  |  | Voice pitch | 0.033 | 0.041 | -0.047, 0.113 | 0.814 | .416 |
| Reproductive domain: moderation analyses of type of masculinity | | | | | | | |
|  |  |  | B | SE | [95% CI] | *z* | *p* |
|  | Masculinity type | **Intercept (body masculinity)** | **0.117** | **0.039** | **0.040, 0.194** | **2.980** | **.003** |
|  |  | 2D:4D | -0.042 | 0.052 | -0.143, 0.059 | -0.812 | .417 |
|  |  | Facial masculinity | -0.030 | 0.071 | -0.170, 0.110 | -0.417 | .677 |
|  |  | **Height** | **-0.107** | **0.038** | **-0.181, -0.033** | **-2.817** | **.005** |
|  |  | Testosterone levels | -0.028 | 0.069 | -0.163, 0.107 | -0.403 | .687 |
|  |  | Voice pitch | -0.032 | 0.079 | -0.187, 0.122 | -0.411 | .681 |
| *Note.* Moderation analyses were only run where each level of the moderator included observations from at least two studies and three independent samples. | | | | | | | |

Supplementary File 6A

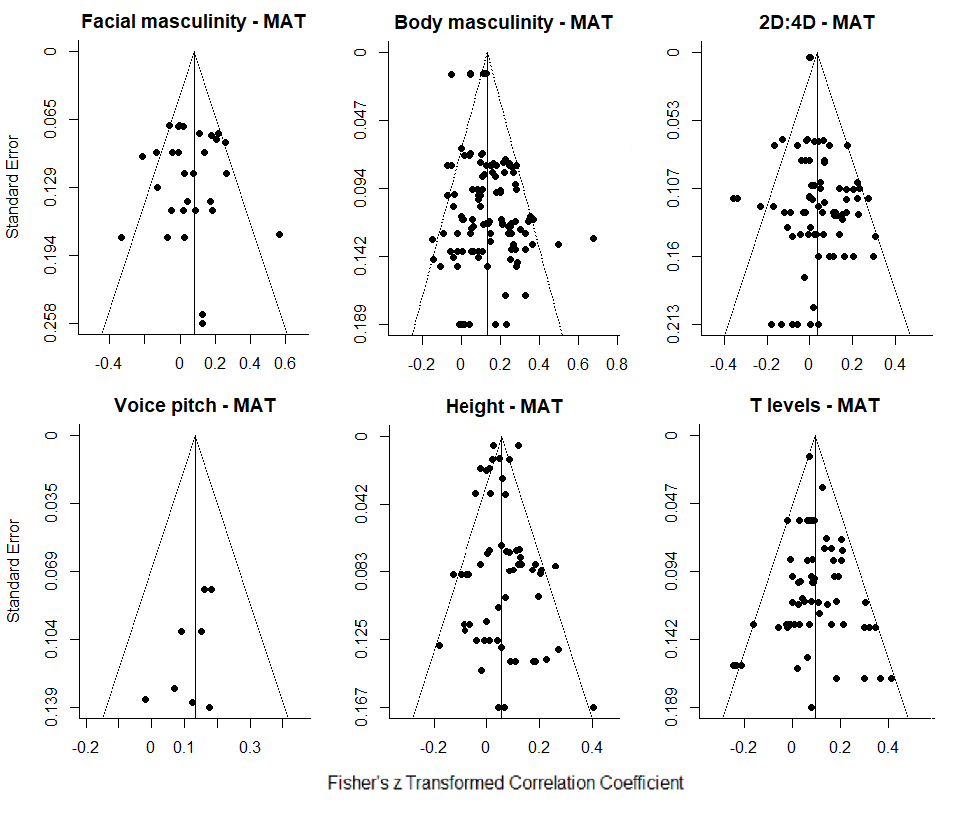
Funnel plots of effect sizes for mating measures (MAT). T = testosterone.

Supplementary File 6B

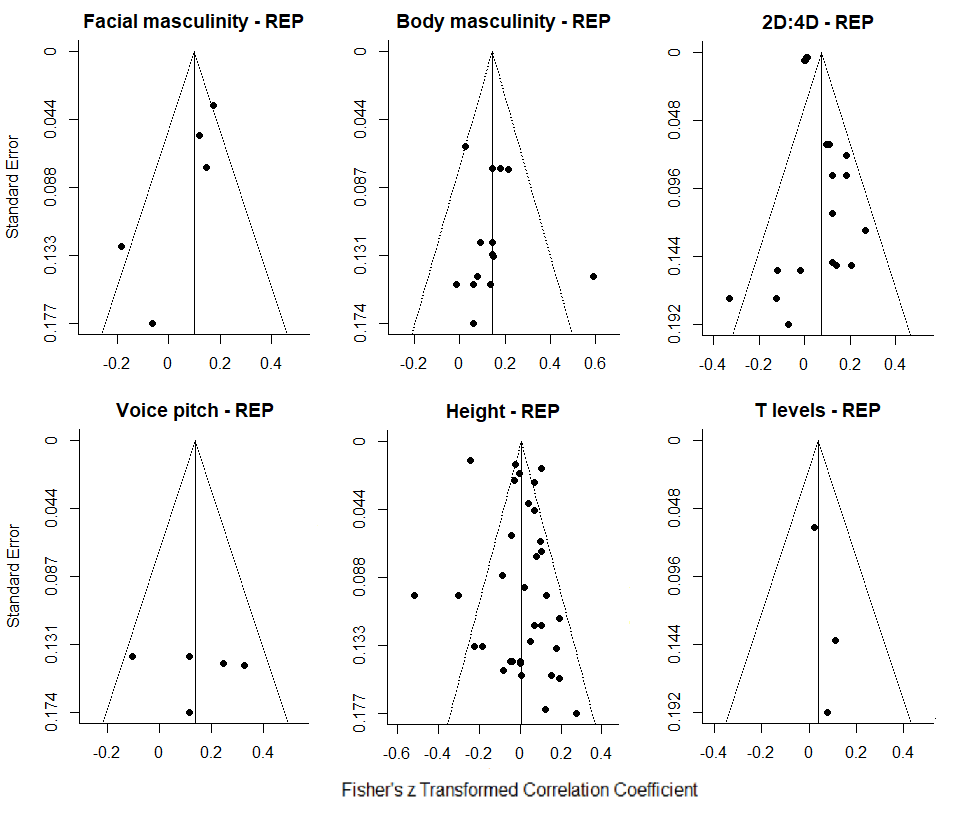

Funnel plots of effect sizes for reproductive measures (REP). T = testosterone levels.

Supplementary File 7

*Output for q-value computation for all analyses*

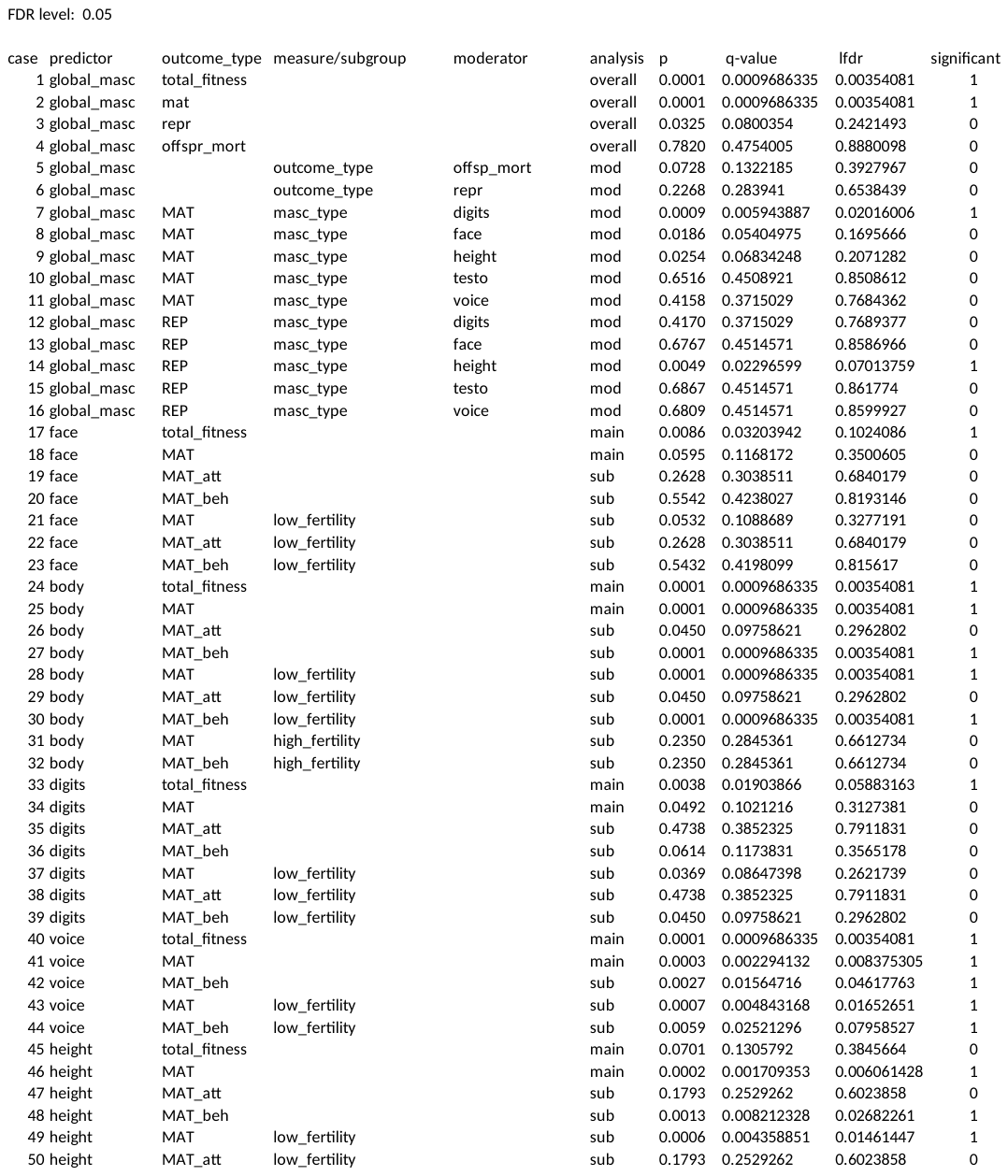

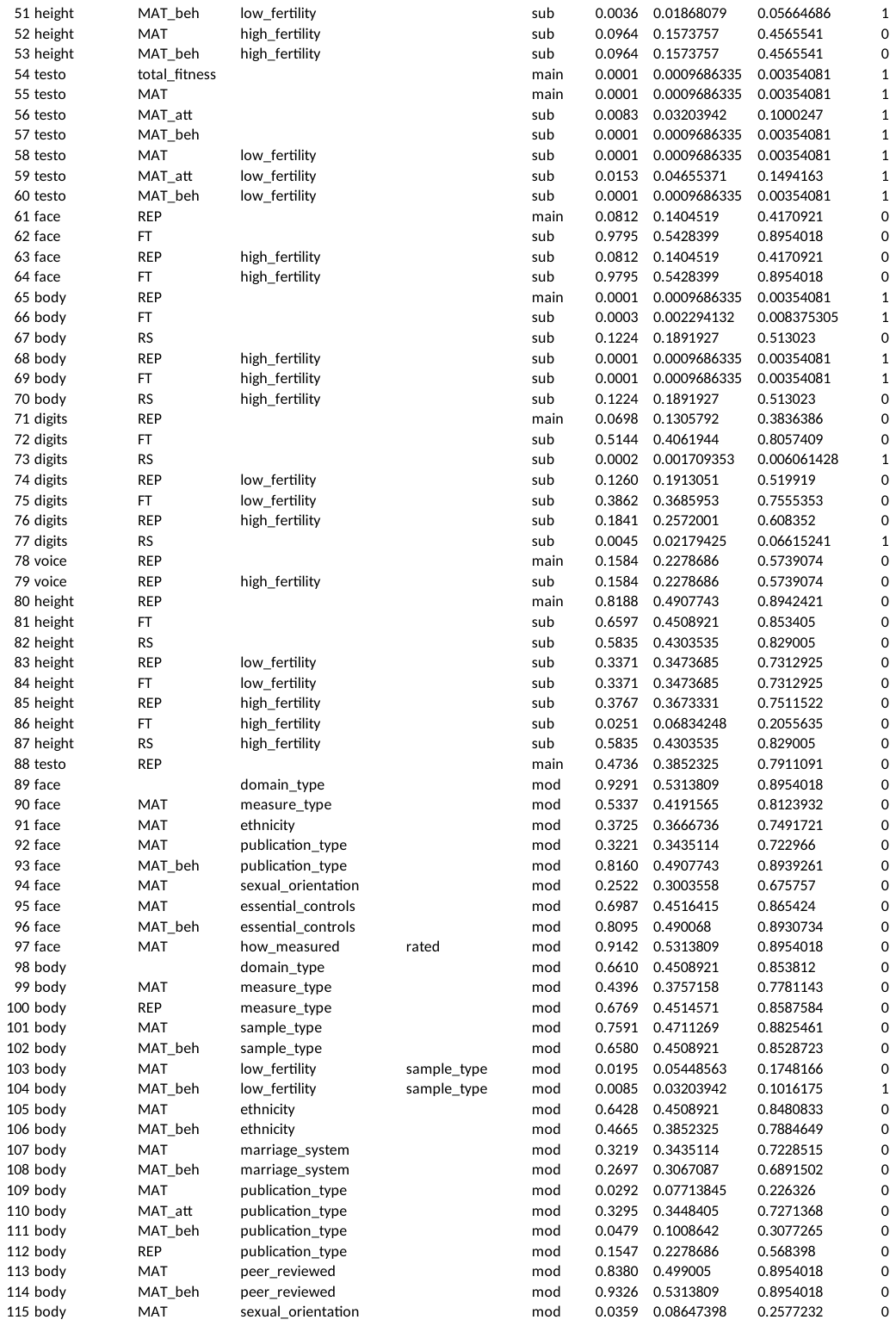

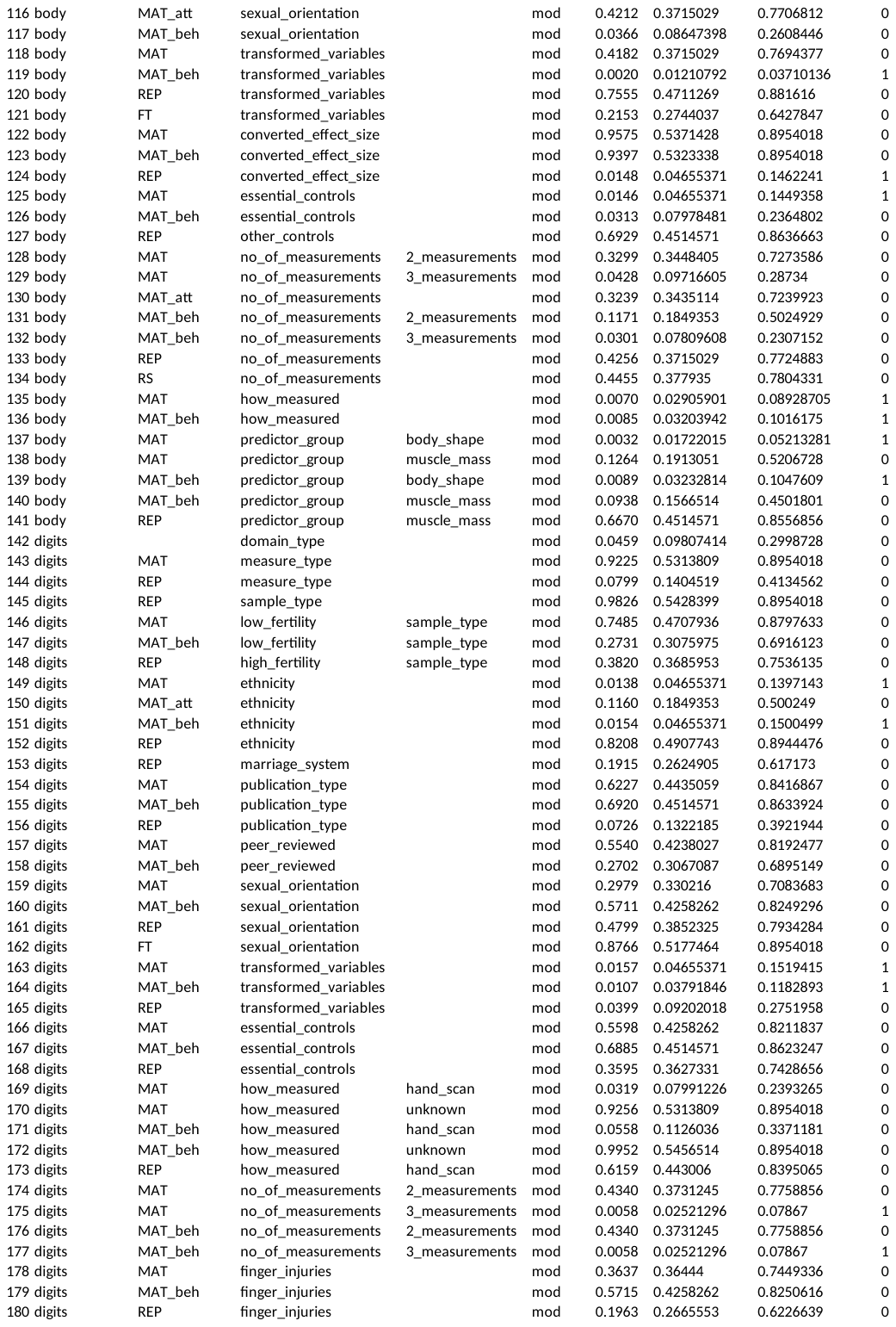

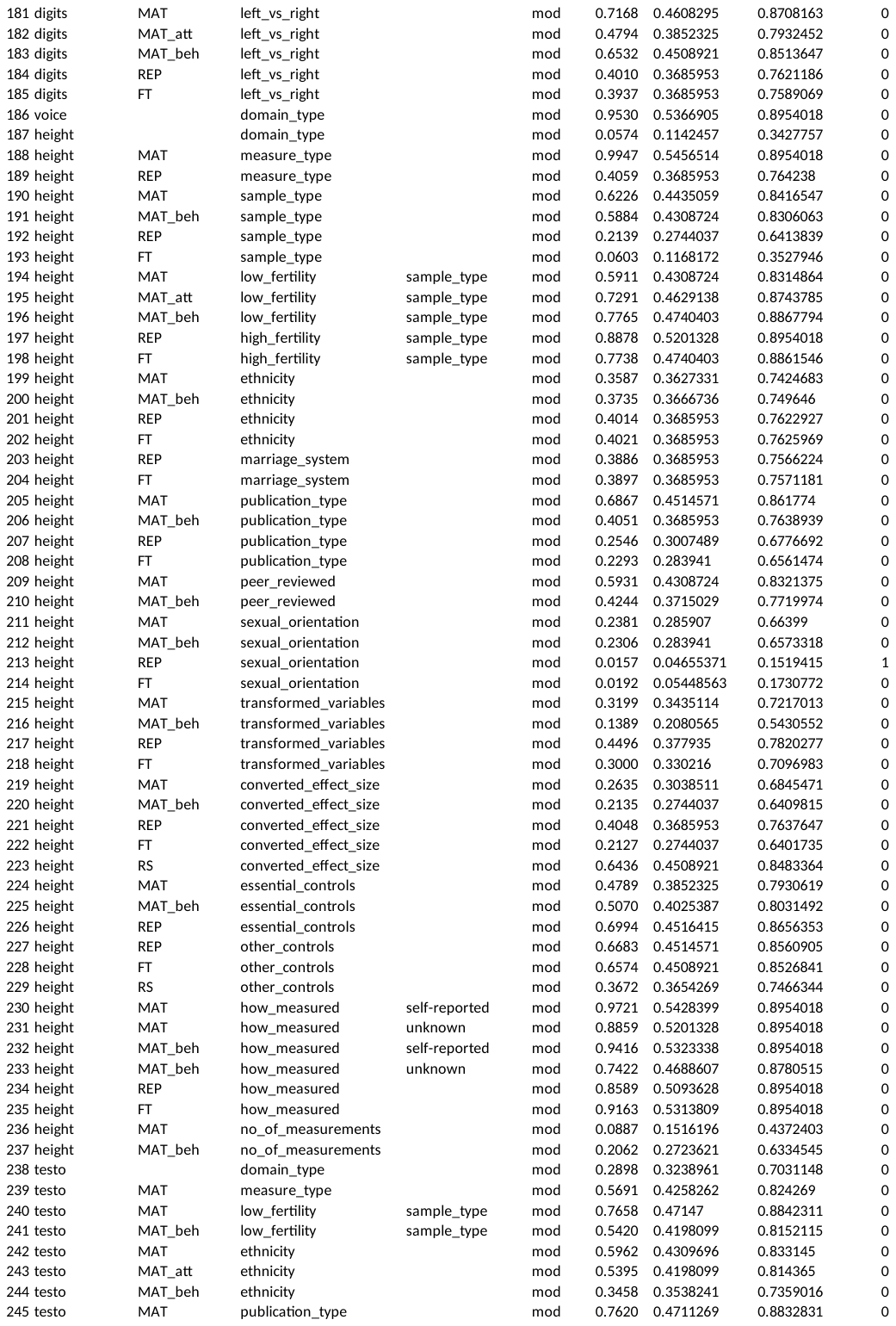

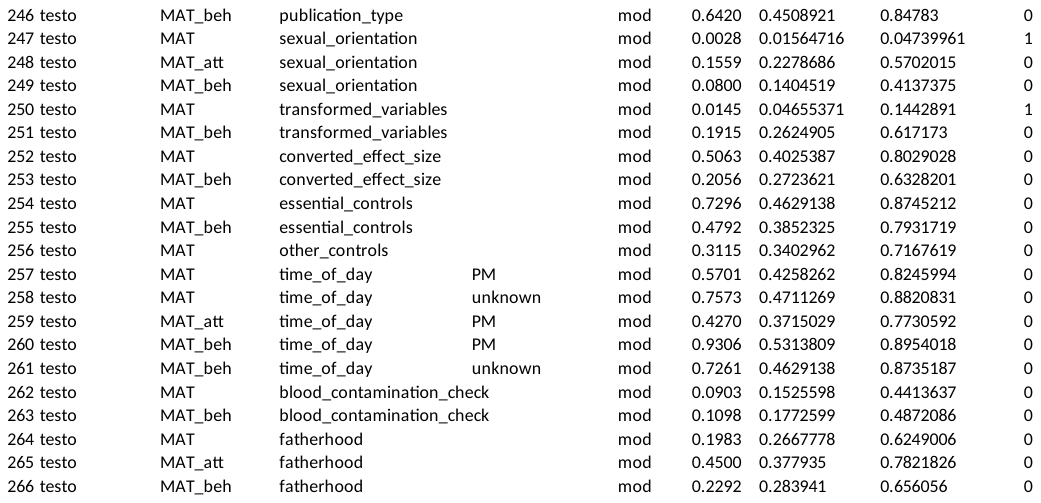
